# Human spinal cord differentiation proceeds rapidly in vitro and only initially maintains differentiation pace in a heterologous environment

**DOI:** 10.1101/2021.02.06.429972

**Authors:** Alwyn Dady, Lindsay Davidson, Pamela A. Halley, Kate G. Storey

## Abstract

Species-specific differentiation pace in in vitro assays indicates that some aspects of neural differentiation are driven by cell-autonomous processes. Here we describe a novel in vitro human neural rosette assay that recapitulates the temporal sequence of dorsal spinal cord differentiation but proceeds more rapidly than in the human embryonic spinal cord, suggesting that in vitro conditions lack endogenous signalling dynamics. To test the extent to which this in vitro assay represents a cell intrinsic differentiation programme, human iPSC-derived neural rosettes were homo-chronically grafted into the faster differentiating chicken embryonic neural tube. Strikingly, in vitro human differentiation pace was not accelerated, even in single host-integrated cells. Moreover, rosette differentiation eventually stalled in a neural progenitor cell state. These findings demonstrate the requirement for timely extrinsic signalling to accurately recapitulate human neural differentiation tempo, and suggest that while intrinsic properties limit differentiation pace, such signals are also required to maintain differentiation progression.

## Introduction

Understanding human embryonic development is a major challenge in contemporary biology and is in large part investigated by creation of pluripotent cell derived in vitro models of developing tissues and organs. One of the most striking features of such in vitro assays is the remarkably faithful recapitulation of specific differentiation programmes in minimal culture conditions. For example, species specific patterns of cell proliferation and timing of neuronal sub-type differentiation are evident in human and macaque cortical progenitors, even when these cells are cultured together (Otani et al. 2016), strongly supporting the notion that cell autonomous mechanisms direct differentiation programmes. Recent analyses of potential mechanisms that account for human specific differentiation tempo have identified general protein stability and cell cycle duration as parameters that constrain this process in spinal cord progenitors (Rayon et al. 2020). Moreover, a human specific gene, *ARHGAP11B*, has recently been shown to promote the enhanced proliferative capacity of human basal cortical progenitors by augmenting a specific metabolic pathway (Namba et al. 2020): and so uncovering a genetic basis for such cell autonomous cell behaviour. Such findings indicate that cell intrinsic mechanisms underpin differentiation programme progression and suggest that this process can proceed from the neural progenitor cell state with minimal extrinsic signals. Evaluation of the extent to which human neural progenitors differentiate cell autonomously requires a detailed understanding of the differentiation timing of specific cell types during normal human embryogenesis and how well this is recapitulated in vitro.

As in other vertebrate embryos, the extracellular environment, including spatially and temporally regulated signals provided by neighbouring tissues, is likely to orchestrate human embryonic differentiation. In vitro models, from rosettes to more complex three-dimensional organoids and gastruloids, aim to recreate this by provision of supportive extracellular environments and timely exposure to ligands or small molecules that modulate signalling activity (Lancaster et al. 2013, Otani et al. 2016, Ogura et al. 2018, Chiaradia and Lancaster 2020, Moris et al. 2020). A critical test of the extent to which neural progenitor differentiation is governed by cell autonomous mechanisms is their behaviour in a heterologous environment. However, studies examining differentiation pace and accuracy in such contexts are lacking. This is obviously important when considering use of such neural progenitors in cell replacement strategies following injury or disease (Sofroniew 2018) or for local delivery of secreted factors for therapeutic effect (Zhang et al. 2017), as these approaches involve transplantation into heterologous environments and yet anticipate differentiation into specific cell types, e.g. (Kumamaru et al. 2018, Kumamaru et al. 2019).

A relatively simple region of the developing nervous system in which to evaluate the extent to which human neural progenitors differentiate cell autonomously, is the developing spinal cord. In vitro generation of spinal cord progenitors from mammalian pluripotent cells has been achieved by approaches in which anterior neural tissue is induced and then caudalised, e.g. (Meinhardt et al. 2014, Gupta et al. 2018). Recent advances in understanding the cellular origins of the spinal cord have further shown that it arises from a bipotent epiblast cell population adjacent to the anterior primitive streak known as neuromesodermal progenitors (NMPs) (Tzouanacou et al. 2009, Gouti et al. 2014, Gouti, Metzis, and Briscoe 2015, Henrique et al. 2015, Gouti et al. 2017, Steventon and Martinez Arias 2017). Moreover, in vitro generation of spinal cord progenitors from NMPs may align better with the temporal sequence of cell fate decisions in the embryo. Derivation of NMP-like cells from pluripotent cells has been demonstrated (Turner et al. 2014, Gouti et al. 2017, Frith et al. 2018, Verrier et al. 2018, Frith and Tsakiridis 2019, Edri et al. 2019) and this has catalysed investigation of mechanisms regulating human as well as mouse spinal cord development.

Here we present a novel human pluripotent cell-derived in vitro neural rosette assay that recapitulates dorsal spinal cord differentiation observed in vertebrate embryos and use this to evaluate to the extent to which human neural progenitors differentiate cell autonomously. Using key cell type-specific marker proteins we compare in detail in vitro neural differentiation with that in the human embryonic spinal cord. In a series of grafting experiments, in which human iPSC derived spinal cord progenitors are transplanted into the chicken embryonic neural tube, we further test whether human differentiation pace can be increased in this faster differentiating environment, as well as the extent to which the differentiation trajectory of transplanted cells relies on timely provision of extrinsic signalling.

## Results

### In vitro generation of human dorsal spinal cord rosettes

An in vitro protocol which generates dorsal spinal cord from human pluripotent cell-derived NMP-like cells with brachial/ thoracic character (Verrier et al. 2018) was used to make neural rosettes (Fig. 1A). The precise timing with which NMP-like cells (derived from H9 hESCs) enter the neural differentiation programme was assessed by quantifying cells expressing a range of neural progenitor markers during the first 10 days of two-dimensional culture in a minimal medium (Figs. 1A,B). This early neural tissue formed compact condensations of cells by day 4 (D4) of NMP-like cell differentiation, which lacked overt apico-basal polarity as indicated by localisation of N-Cadherin (Cadherin-2) at most cell-cell interfaces (Hatta and Takeichi 1986, Dady, Blavet, and Duband 2012) (Fig. 1B). These cells expressed SOX2, characteristic of both pluripotent, NMP and neural progenitor cells (Uchikawa et al. 2003, Uchikawa et al. 2011, Bylund et al. 2003) (Ellis et al. 2004), and the neural progenitor protein PAX6 (Walther and Gruss 1991, Pevny et al. 1998). Some cells expressing the early neural crest marker SNAI2 were also detected on D4 (Nieto 2002, Jiang et al. 2009, Dady and Duband 2017) (Fig. 1B). The expression of PAX7, which distinguishes dorsal neural progenitors (Jostes, Walther, and Gruss 1990) (Gruss and Walther 1992) and the absence of OLIG2, characteristic of more ventral spinal cord (Mizuguchi et al. 2001, Novitch, Chen, and Jessell 2001) further indicated that at D4 this differentiation protocol had generated human neural tissue with dorsal spinal cord identity (Fig. 1B).

**Figure 1.**
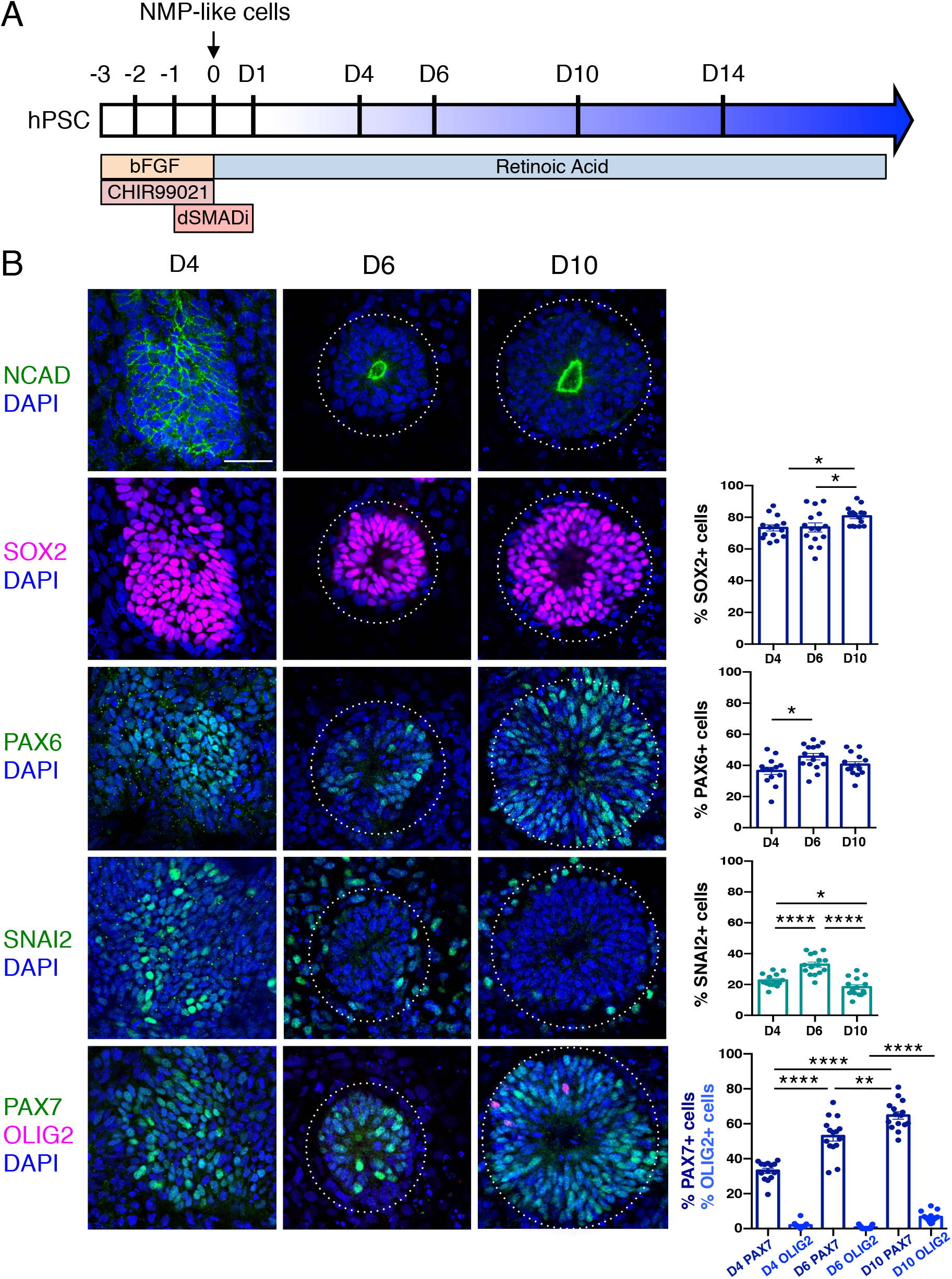
Human pluripotent cell derivation of NMP-like cells and their differentiation into dorsal spinal cord neural rosettes. (**A**) Timeline and protocol for induction and differentiation of dorsal spinal cord neural rosette from human pluripotent stem cells (hPSC); (**B**) Expression patterns and quantification of cell type specific proteins in rosettes on days (D)4, D6 and D10 detected by immunofluorescence combined with heterochromatin marker DAPI: apical polarity marker N-Cadherin (NCAD), neural progenitor markers SOX2, PAX6, early neural crest marker SNAI2, and dorso-ventrally restricted neural progenitor markers PAX7 and OLIG2. Quantifications indicate proportions of expressing cells in either a defined field on D4 or within a rosette D6 and D10, in 15 samples for each marker (n=5 from each of 3 independent differentiations, see Methods). Data analyzed with Mann-Whitney test, errors bars ± S.E.M, p-values = * p < 0.05, ** p < 0.01 and **** p < 0.0001. Scale bar = 50μm.

By D6 neural rosettes composed of apico-basally polarised SOX2, PAX6 and PAX7 expressing progenitors had formed, with a single, central lumen and peripheral SNAI2 neural crest cells (Fig. 1B). This spatial reorganisation was accompanied by decline in SNAI2+ cells and the appearance, by D10, of a scattering of OLIG2 expressing cells (Fig. 1B). This emergence of neural rosettes from a condensing epiblast cell population at D4, may equate to the formation of the neural plate and establishment of the pseudostratified neuroepithelium, but it is also reminiscent of the phases of embryonic secondary neurulation (Schoenwolf and Nichols 1984, Catala, Teillet, and Le Douarin 1995, Dady, Blavet, and Duband 2012, Dady et al. 2014, Fedorova et al. 2019).

These findings show that differentiation of human ESC derived NMPs in a minimal culture medium containing retinoic acid (RA) is sufficient to generate self-organising dorsal spinal cord rosettes and accompanying neural crest. Moreover, this protocol induced similar dorsal spinal cord differentiation in hiPSC-derived NMP-like cells (Fig. S2).

### Sequential differentiation of human dorsal spinal cord progenitors is recapitulated in vitro

Dorsal spinal cord rosettes were further characterized as they progressed through their differentiation programmes (Fig. 2A). Decline of SNAI2-expressing cells at D10 coincided with the emergence of migrating neural crest cells expressing HNK1 (Tucker et al. 1984, Bronner-Fraser 1986), SOX10 (Bondurand et al. 1998) and TFAP2α (Zhang et al. 1996, Betters et al. 2010) in cells now just outside the rosettes (Jiang et al. 2009) (Fig. 2B). The onset of neuron production was assessed in several ways; (i) by analysis of immature neuronal marker Doublecortin (DCX) expression (Gleeson et al. 1999, Brown et al. 2003) (using a hESC (H1) neuronal reporter cell line in which a yellow fluorescent protein (*YFP*) sequence is fused to endogenous *DCX* (Yao et al. 2017) (Fig. 2C), and ii) by characterization of cdk inhibitor P27/KIP1 expression, which identifies post-mitotic cells in the neural tube (Gui, Li, and Matise 2007) (Fig. 2D). These analyses defined a neurogenesis timeline which commenced by D8, peaked at D14 and progressed through to D20.

**Figure 2.**
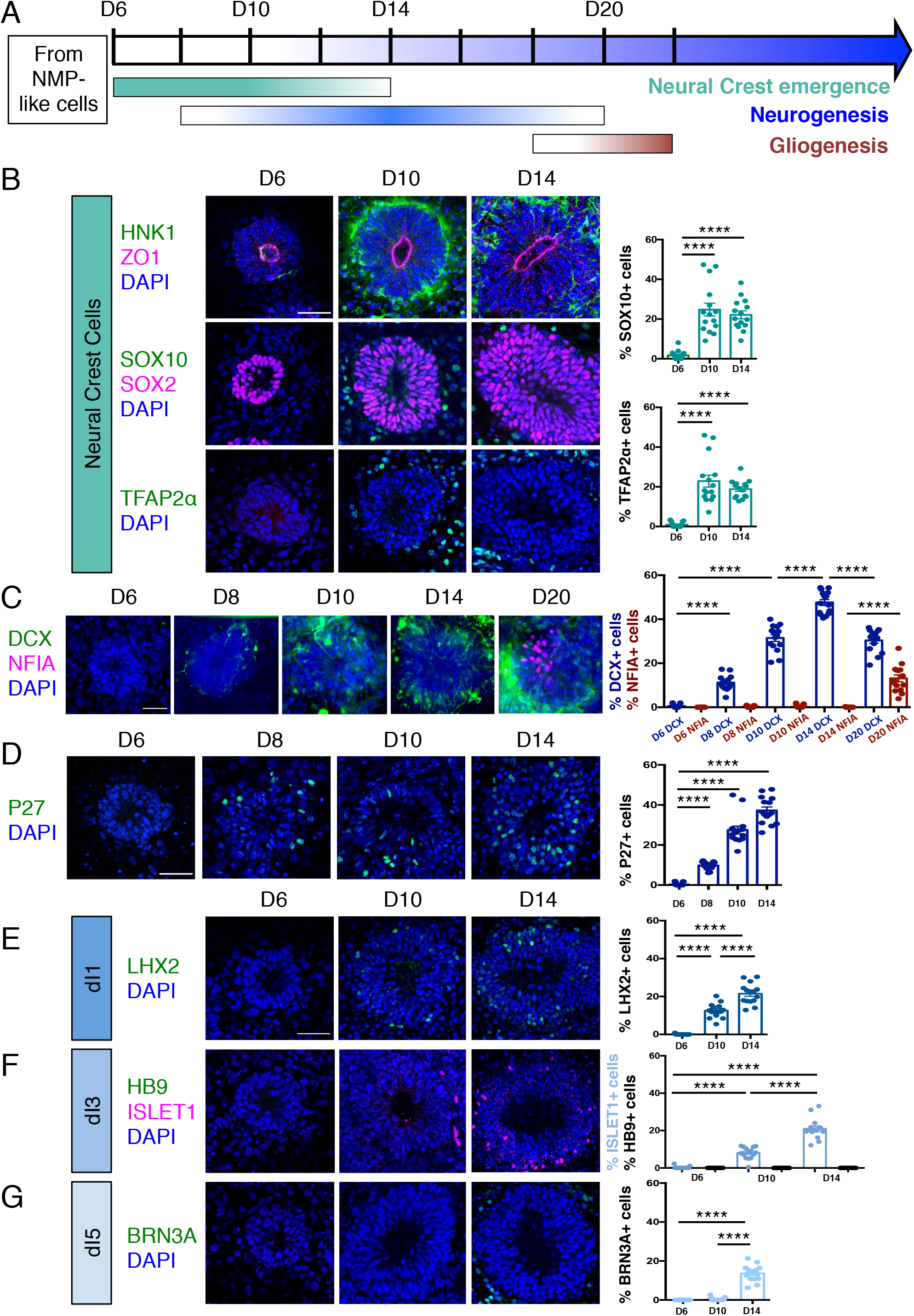
Human dorsal spinal cord rosettes exhibit stereotyped sequence of cell-type specific differentiation. (**A**) Schematic of in vitro sequence of cell-type specific differentiation of dorsal spinal cord rosettes; (**B**) Emergence of migrating neural crest from rosettes by D10, documented by IF for key marker proteins HNK1, (lumen defined by apical marker ZO-1), SOX10 (neural progenitors defined by SOX2) and TFAP2α and quantified for SOX10 and TFAP2α; (**C**) Transition from neurogenesis to gliogenesis documented and quantified in immature neuronal reporter line (H1:DCX-YFP) with co-IF for gliogenesis marker Nuclear Factor I-A (NFIA) at key timepoints; (**D**) P27/KIP1 expression which by analogy identifies most dorsal post-mitotic neurons, confirms neurogenesis onset by D8; (**E-G**) Emergence of dorsal interneuron (dIs) subtypes by D10 indicated by dI1s/LHX2 (**E**), dI3s/ISLET1 (**F**) (note lack of HB9/ ISLET1 co-expression distinguishes these cells from motor neurons) and dI5s/BRN3A (**G**). Sample numbers, statistics and error bars as in Figure 1, each dot represents a single rosette and see Materials and Methods; Scale bar = 50μm.

By D20 DCX-expressing neurons largely resided at the rosette perimeter and the early glial progenitor marker, the transcription factor Nuclear factor 1 A-type (NFIA) (Deneen et al. 2006), was now detected in cells within the rosette (Fig. 2C). This indicates operation of a switch from neurogenic to gliogenic differentiation programmes in the human spinal cord rosettes that is similar to that observed in the mouse embryo (Deneen et al. 2006).

Finally, we determined the identity of neurons generated in this in vitro assay by immunofluorescence of dorsal interneuron (dIs) subtype-specific markers. In the mouse embryonic neural tube distinct sensory interneurons arise in specific dorsoventral positions and express distinguishing marker proteins (Lai, Seal, and Johnson 2016). By D10, proprioceptive interneurons (dI1s) and mechanosensory interneurons (dI3s) were detected by the expression of LHX2 (Fig. 2E) and ISLET1 (Fig. 2F), respectively (Liem, Tremml, and Jessell 1997). By D14, dI5 interneurons expressing BRN3A were detected (Ninkina et al. 1993, Fedtsova and Turner 1995) (Fig. 2G). Moreover, ISLET1+ cells did not co-express the transcription factor HB9, a marker of spinal cord motoneurons (Arber et al. 1999), confirming their dI3 identity (Fig. 2F). In contrast, LHX1/dI2s/dI4s expressing cells were not identified (Fig. S1), indicating that this protocol is insufficient to generate all types of dorsal interneurons.

These data document the differentiation capacity in this human spinal cord rosette assay and demonstrate that it reproducibly generates dorsal neural differentiation programmes. These begin with dorsal neural progenitors and emerging neural crest, progress to neurogenesis (of specific dorsal interneurons dI1s, dI3s and dI5s) and a later switch to gliogenesis: a sequence that recapitulates the temporal order of differentiation observed in the spinal cord of mouse and avian embryos (Fig. 2A) (Deneen et al. 2006, Glasgow et al. 2017, Andrews et al. 2017).

### Human spinal cord differentiation in vitro progresses more rapidly than in the human embryonic spinal cord

This conservation in the sequence of the dorsal neural differentiation programme contrasts with the longer duration of human embryogenesis in comparison with that observed in the mouse and chicken. Indeed, the pace of differentiation in the developing nervous system and that of body axis segmentation is slower in human embryos (Rayon et al. 2020, Matsuda et al. 2020). These latter studies suggest the presence of species-specific cell intrinsic differences that govern developmental tempo. To understand how well this tempo is captured in our in vitro assay we next sought to compare the sequence and timing of differentiation progression in human spinal cord rosettes with that observed in the human embryo.

The pattern and timing of key marker protein expression in human embryonic spinal cord was assessed using a combination of tissue samples provided by the HDBR (Figs. 3A-H) and available data. One way in which to assess differentiation progression is to take advantage of the spatial separation of the temporal events of differentiation along the embryonic body axis, and so analyses were carried out at distinct rostro-caudal levels in Carnegie stage (CS) 12/13 (gestational weeks 4-4.5) human embryos (O’Rahilly 1987); sacral (the most caudal and least differentiated region), lumbar and thoracic/brachial (more rostral and most differentiated region). Developmental time along the body axis can be approximated by that indicated by segmentation progression from the sacral to thoracic region, which involves generation of ∼20 somites (with 1 human somite generated every 5 hours (Turnpenny et al. 2007, Hubaud and Pourquié 2014, Matsuda et al. 2020) this represents, 100h or ∼4 days (Fig. 3A).

**Figure 3.**
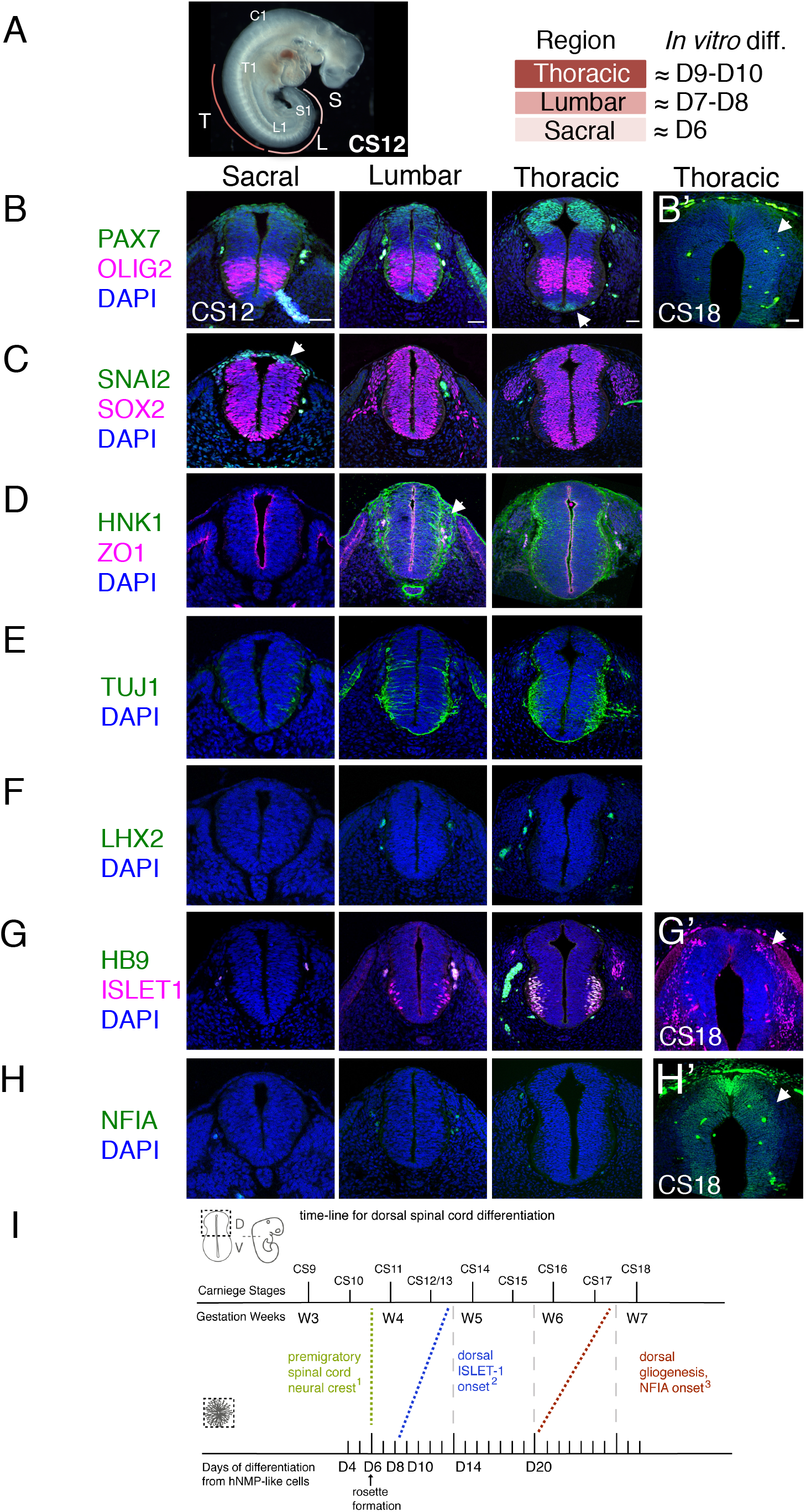
Characterization of neural progenitors and neuronal subtypes in human embryonic spinal cord and comparison with in vitro neural rosette differentiation. (**A**) CS12 human embryo with anatomical sub-regions indicated and their alignment with in vitro spinal cord rosette differentiation (see text); (**B-H**) transverse sections through human embryonic spinal cord at sacral, lumbar and thoracic levels at 4/4.5 weeks (CS12/CS13) and at brachial/thoracic at 7/7.5 weeks (CS18) analysed by IF for expression of (**B and B’**) dorsal, PAX7, or ventral, OLIG2, spinal cord progenitor markers (note novel PAX7 detection in the floor plate, arrow) and **(B’**) waning PAX7 at CS18 (arrow); (**C**) premigratory neural crest cell marker SNAI2 is only detected in sacral regions (arrow) shown here with pan-neural progenitor marker SOX2; (**D**) differentiating migrating neural crest expressing HNK1 emerges from the lumbar region (arrow); (**E**) neuronal differentiation detected with pan-neuronal marker TUJ1 is manifest dorsally in lumbar regions; dorsal interneurons indicated by (**F**) dI1 specific transcription factor LHX2 and (**G**) dI3 identifying ISLET1 expressing cells were not detected dorsally in any region of the spinal cord at CS12/CS13; (G’) dorsal ISLET1 interneurons at CS18; (**H**, **H’**) gliogenesis marker NFIA detected dorsally only at CS18; (**I**) Schematic aligning dorsal brachial/thoracic spinal cord differentiation in human embryo and in hPSC derived dorsal spinal cord rosettes, based on (1) Bondurand N. et al. 1998, O’Rahilly and Müller 2007; (2) This study, Rayon T. et al. 2020, Marklund et al 2014; (3) This study, Rayon T. et al. 2020); three sections were analysed at each level for each marker in n = 2 CS12/13 and n = 1 CS18 human embryos. Scale bar = 50μm.

**Figure S1.**
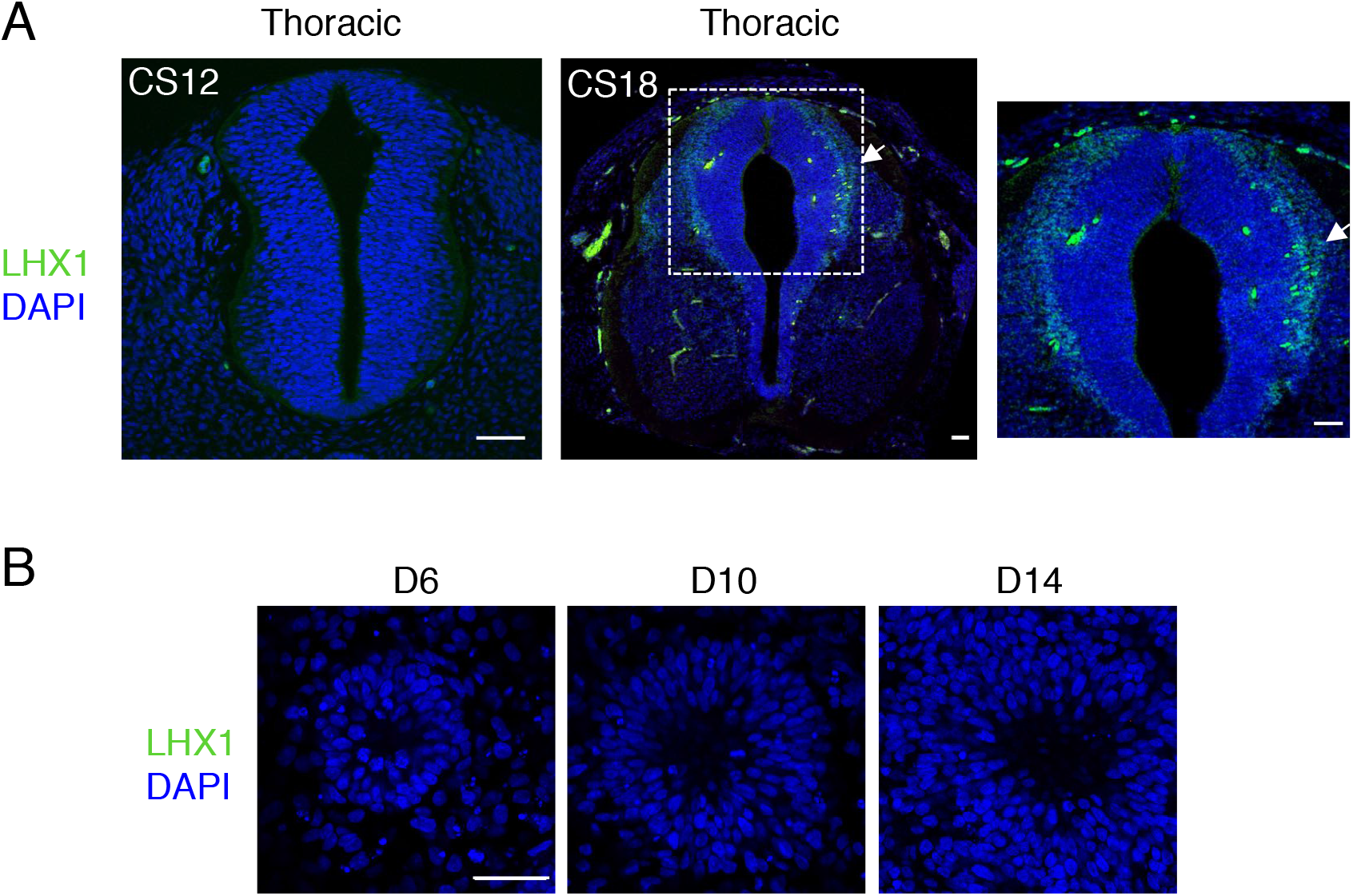
Dorsal interneurons type 2 expressing LHX1, dI2/LHX1, are not generated in human spinal cord neural rosettes in vitro. (**A**) dl2 neuron differentiation analysed by IF for expression of LHX1 in stages CS12 and CS18 human embryonic spinal cord; note dI2 neurons are only detected at CS18 (arrow indicates LHX1 expressing cells, dashed white boxes indicate area of magnification in next adjacent image); (**B**) LHX1 is not detected in neural rosettes differentiated in vitro. Three sections were analysed at each stage in n = 1 CS12 n=1 CS13 and n =1 CS18 human embryos. Three independent cell differentiations were performed to generate rosettes. Five neural rosettes from each experiment were then analysed for quantification at each time point (n=15). Data were analysed using non-parametric Mann-Whitney test. Scale bar = 50μm

The sacral spinal cord at CS12/13 stage was composed of progenitors expressing dorsal (PAX7) and ventral (OLIG2) markers (Fig. 3B) as well as pre-migratory neural crest cells expressing SNAI2 (Fig. 3C) and lacking migratory neural crest cell marker HNK1 (Fig. 3D). Moreover, neurogenesis, revealed by the pan-neuronal marker TUJ1, was only detected ventrally (Menezes and Luskin 1994) (Figs. 3E). The differentiation state in the dorsal region of the CS12/13 sacral neural tube therefore aligns well with that found in D6 dorsal spinal cord rosettes, which constitute apico-basally polarised neuroepithelium expressing PAX6, PAX7 and SNAI2 (Fig. 1B) that lacks migratory neural crest and neurons (Figs 2B-G).

In the more rostral, lumbar region, PAX7 and OLIG2 domains have enlarged (Marklund et al. 2014) and, while SNAI2-expression has declined (Fig. 3C), migrating HNK1-expressing neural crest cells have now emerged (Fig. 3D). The first neurons are now detected in the dorsal half of the neural tube as indicated by TUJ1 expression (while co-expression of ISLET1 and HB9 identifies ventrally located motor-neurons (Arber et al. 1999, Rayon et al. 2020), Figs. 3E, G). The dorsal neurons may correspond to the DCX /P27 expressing cells detected in D8 rosettes (Figs. 2C,D), however, their identity remains undefined. Neurons in this dorsoventral position (ventral region of dorsal half of the neural tube) in the mouse and chicken neural tube express LHX1 (Lai, Seal, and Johnson 2016, Andrews et al. 2017), but this protein was not detected in neural rosettes and appears much later in the human embryo (Fig. S1). Given the alteration in transcription factor combinatorial code created by the expansion of the OLIG2 domain in human spinal cord progenitors (Marklund et al. 2014), one possibility is that human-specific neuronal cell types arise in this region. The dorsal region of the lumbar neural tube therefore broadly aligns with D8-D10 dorsal spinal cord rosettes in which neuronal differentiation and neural crest migration have now commenced (Figs. 2B-D).

Importantly, dorsal interneurons dI1 and dI3 indicated by LHX2 and ISLET1/HB9-expression respectively were not detected in the human CS12 or CS13 embryonic spinal cord at lumbar or the thoracic levels assessed here (which included sections from presumptive forelimb level, brachial spinal cord), (Figs. 3F, G), although a few dorsal ISLET1 positive cells were detected in brachial spinal cord of a further CS13 embryo (T. Rayon pers comm.). This suggests that these interneurons are just beginning to be generated at CS13. In contrast, ISLET1 and LHX2 positive cells were found in all D10 neural rosettes (Figs. 2E, F). This suggests that the dorsal interneuron differentiation programme progresses more rapidly in vitro than in the human embryonic spinal cord.

A further way to assess differentiation tempo is to align progression of a defined cell population in vitro and in vivo using time elapsed from a key landmark event. The NMP-like cells generated in our protocol have a brachial/thoracic regional character as indicated by Hox gene expression (Verrier et al. 2018) and differentiation of this region can be compared in vitro and in embryos over time. Detailed analysis of the first appearance of spinal cord neural crest in the human embryo provides a clear landmark with which to align in vivo and in vitro differentiation (Bondurand et al. 1998, O’Rahilly and Muller 2007). These latter studies identify morphological emergence of migrating neural crest in cervical level spinal cord at CS11 human embryo (O’Rahilly and Muller 2007) and in even more caudal spinal cord at this stage using *SOX10 mRNA* expression (Bondurand et al. 1998). From this we infer that pre-migratory neural crest is present prior to CS11 in brachial/thoracic spinal cord, and so aligns this region in CS10-11 embryos with D6 in our neural rosettes, which express SNAI2 but lack migratory neural crest markers proteins SOX10, HNK1 and TFAP2α (Figure 1B and 2B). Using this alignment point we then compared time to appearance of later cell type specific markers in neural rosettes and human embryos (Fig.3I). As above, dorsally located ISLET1/dI3s first appear ∼ CS13 and are robustly detected at CS15 ((Rayon et al. 2020); and see HDBR Atlas http://hdbratlas.org), and here this indicates that these neurons appear at least ∼4 days earlier in D10 rosettes (Figs. 2E,F, 3I). The landmark switch from neurogenesis to gliogenesis indicated by NFIA expression in dorsal neural progenitors was detected in human embryos at CS18 (gestational weeks 7-7.5) (Fig. 3H’) and not at CS17 (Rayon et al. 2020). This is ∼ 1 week later than in neural rosettes, where NFIA was robustly found at D20 (Figs. 2C, 3I). Moreover, the overall time from spinal cord migratory neural crest appearance to dorsal gliogenesis takes ∼ 2 weeks (D6 -D20) in human dorsal spinal cord rosettes *in vitro* and ∼ 3-3.5 weeks (CS10/11-CS17/18) in the human embryonic spinal cord (Fig. 3I). These findings indicate that in vitro conditions can alter the pace of neural differentiation progression, which proceeds faster than in vivo within the context of these self-organizing neural rosettes.

### Cell intrinsic differentiation of human spinal cord progenitors is unaltered by differentiation cues from the more rapidly developing chicken embryonic environment

In the vertebrate embryo, the timing of neural differentiation is orchestrated by signals from adjacent tissues, which drive and coordinate axial differentiation (Diez del Corral, Breitkreuz, and Storey 2002, Diez del Corral et al. 2003) but the localised and temporally controlled characteristics of this regulatory influence are lacking in vitro. Indeed, in the embryo, neural progenitors will generate more cells to make a bigger structure and so proliferative phases that expand cell populations might be longer. We have found that the smaller rosette structures generated in vitro differentiate more rapidly and this raises the possibility that differentiation pace in this minimal assay reflects the species-specific intrinsic constraints on this process (Rayon et al. 2020). To test whether rosette cells retain such a cell intrinsic differentiation programme or if this can run still faster in a more rapidly developing embryo that also provides a more coherent extracellular environment, we next homo-chronically grafted human neural rosettes into the dorsal neural tube of the chicken embryo.

To this end, human iPSCs (Cellartis hIPS4 cell line engineered to express a plasma membrane tagged GFP, ChiPS4-pmGFP, see Methods) were differentiated into NMPs and then into D6 neural rosettes as described above (Fig. 1A). Multiple plates of rosettes were generated so that control in vitro differentiation of iPSC derived rosettes was monitored in parallel with that of D6 rosettes grafted into the chicken embryo. Expression of key marker proteins in these iPSC derived rosettes was found to be the same as that in D6 rosettes derived from the hESC H9 line (Fig. S2). Individual neural rosettes were micro-dissected, lightly dissociated and grafted in place of one side of the dorsal neural tube adjacent to newly formed somites in stage HH10-11 (Embryonic day 2, E2) chicken embryos (Figs. 4A, S3). At this anatomical level the host chicken dorsal neural tube and grafted human D6 rosette are composed of similar populations of dorsal neural progenitors and neural crest expressing respectively PAX7 and SNAI2 and lack neurons and migrating neural crest (Figs. S2, S3); moreover, the D6 rosettes which have brachial/thoracic regional character where grafted into the corresponding region in the developing chicken embryo (opposite somites 10 - 16) providing a regionally appropriate homochronic context in which evaluate differentiation progression. After two days incubation, grafted cells retained apico-basal polarity as indicated by localized ZO-1 and N-cadherin expression (Aaku-Saraste, Hellwig, and Huttner 1996, Hatta and Takeichi 1986, Dady, Blavet, and Duband 2012) (Fig. 4C) and the majority of cells had formed rosette-like structures, which were adjacent but contiguous with the host neuroepithelium, while subsets of cells were incorporated within the host neuroepithelium, including some single cells (Figs. 4B, B’). At this time, most grafted cells still expressed neural progenitor proteins (SOX2 and PAX7)(Figs. 4D, 4E) and the first P27 expressing neurons appeared (Figs 4F and S2), but ISLET-1 expressing (dI3) interneurons were not yet detected (Fig. 4G). This is in good alignment with neural differentiation progression observed in the now D8 rosettes cultured in parallel in vitro (Fig. S2). Importantly, at this stage in the chicken embryo ISLET-1/dl3 expressing neurons are detected (Fig. 4G) (Andrews et al. 2017), consistent with the faster differentiation of the host tissue and indicating that the pace of neural differentiation in grafted human rosettes is unaltered in this more rapidly differentiating environment.

**Figure 4.**
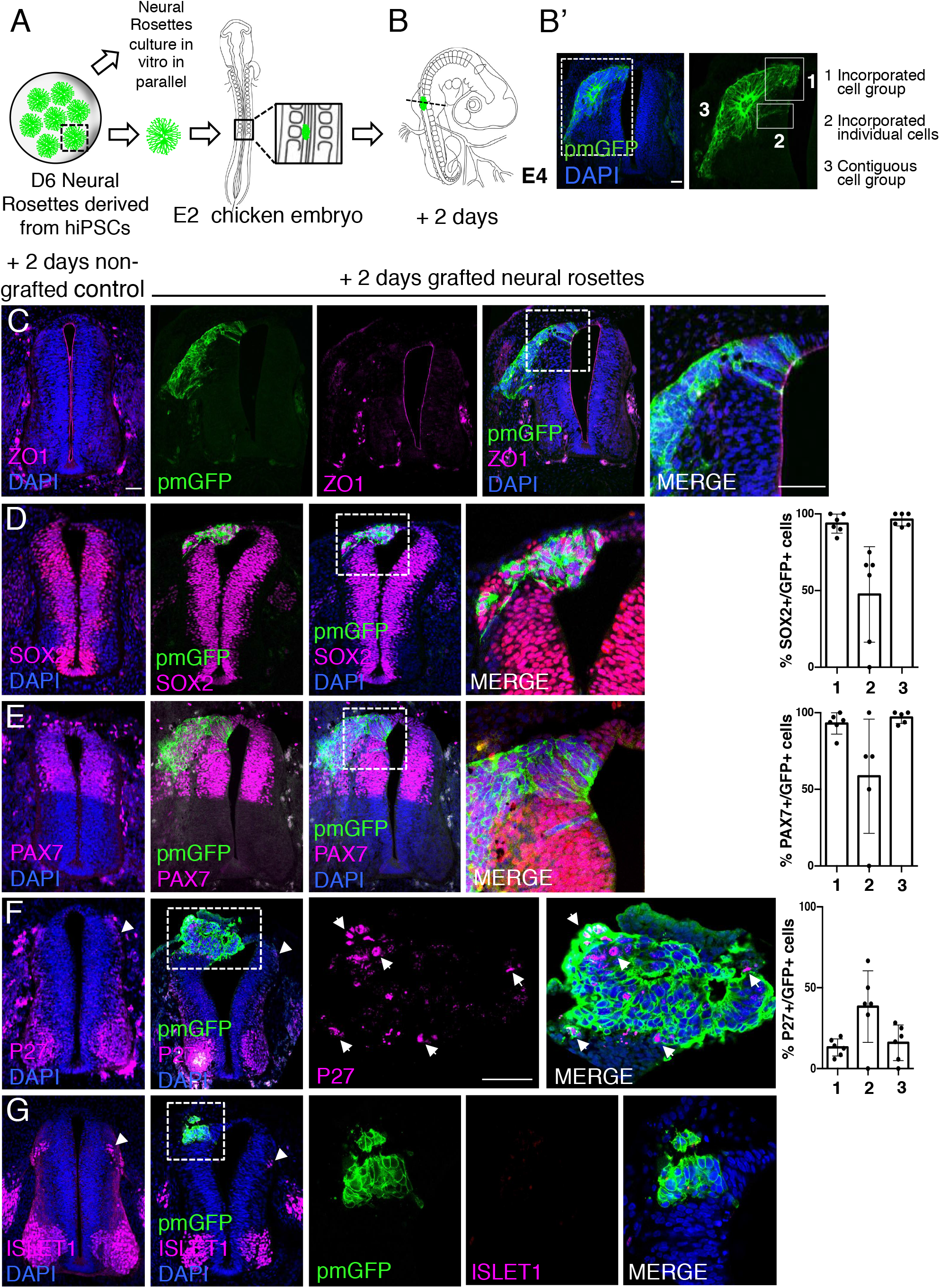
Homo-chronically transplanted hiPSC-derived neural rosettes exhibit cell autonomous differentiation despite exposure to more rapidly developing chicken embryonic environment. (**A**) Schematic of human dorsal spinal cord neural rosette transplantation experiment; (**B-B’**) Two days after grafting human cells were observed in: 1) cell groups or 2) isolated individual cells incorporated within the chicken neural tube, or 3) rosette-like configurations contiguous with chick neural tube. (**C-G**) IF to detect key marker proteins in control (non-grafted, note these sham operated embryos regenerate the dorsal neural tube by 2 days post-operation) and grafted embryos after 2 days incubation (**C**) ZO1, (**D**) SOX2, (**E**) PAX7, (**F**) P27, (**G**) ISLET1 (arrowheads indicate examples of candidate protein expressing cells in the chicken host tissue; arrow indicate example of P27 and GFP co-expressing cells, dashed white boxes indicate area of magnification in next adjacent image), three transverse sections from each of at least three grafted chicken embryos were analysed for each marker combination and the proportions of SOX2/GFP +ve, PAX7/GFP +ve and P27/GFP +ve cells in configurations 1, 2 or 3 were quantified, each dot represents data from one histological section, note the small number of isolated cells (group 2) underlies lack of significant difference between groups (see Materials and Methods), data analysed with Mann-Whitney U test with. Errors bars are ± SD. Scale bar = 50μm.

**Figure S2.**
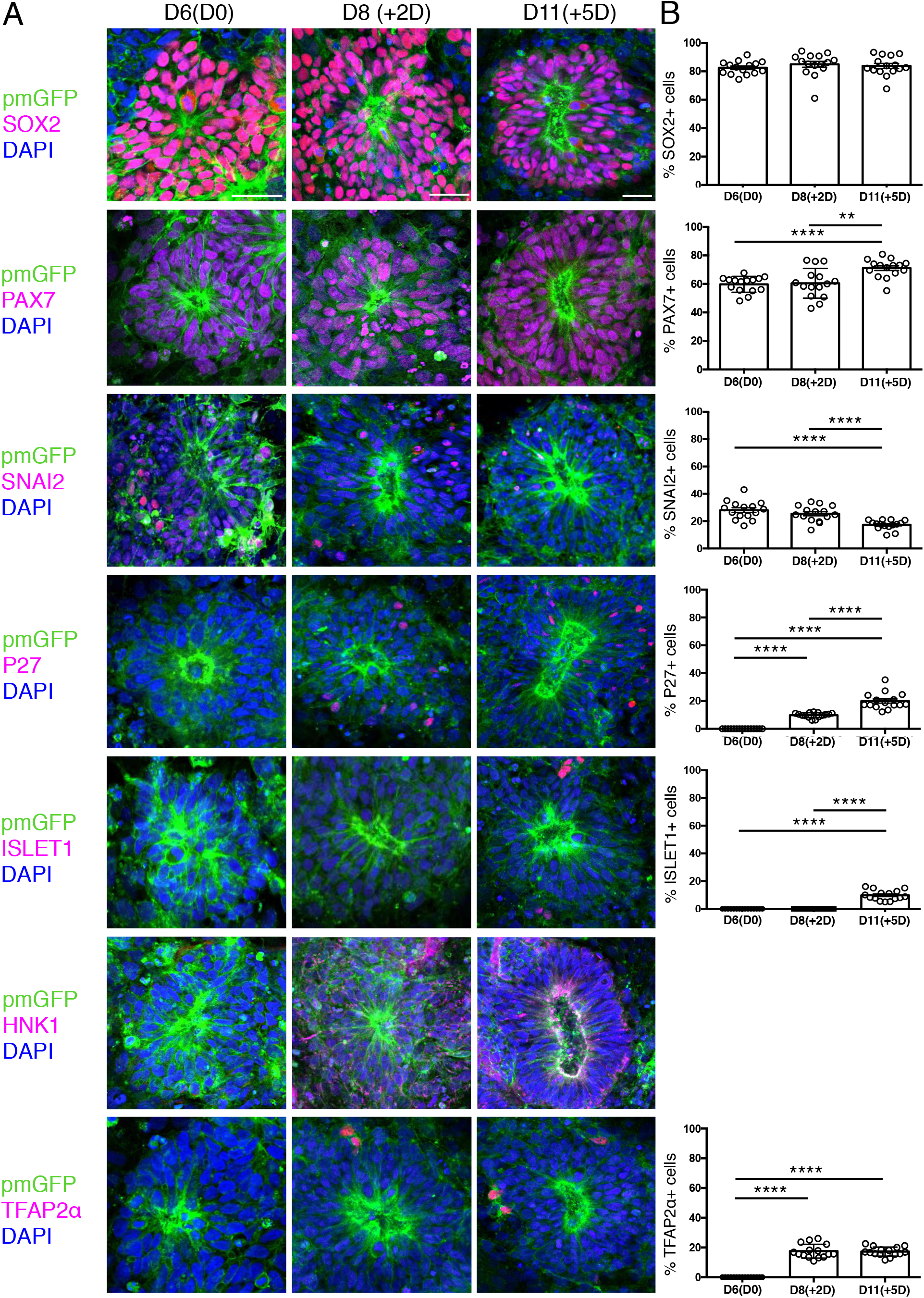
Expression patterns of key marker proteins in human dorsal spinal cord rosettes cultured in vitro in parallel with those homochronically grafted into chick spinal cord. (**A**) Human dorsal spinal cord rosettes from the same differentiation assays used for homochronic grafting experiments were cultured in parallel in vitro and subjected to IF for SOX2, PAX7, SNAI2, P27, ISLET1, HNK1 and TFAP2α SOX2, PAX7 and SNAI2 were expressed from D6(D0). P27, HNK1 and TFAP2α were detected from D8(+2D). ISLET1 was detected from D11(+5D). (**B**) Quantifications, percentage of cells in D6 rosettes expressing each marker on the day of grafting operation D6(D0), two Days after transplantation (+2D) which equals D8 in vitro, and five Days after transplantation (+5D), which equals D11 in vitro differentiation. Five rosettes from each of 3 independent differentiation experiments were used for quantifications, each dot represents data from a single rosette. See methods for statistical analysis. Errors bars are ± S.E.M. Probability of similarity, p-value ** p < 0.01 and **** p < 0.0001. Scale bar = 50μm.

**Figure S3.**
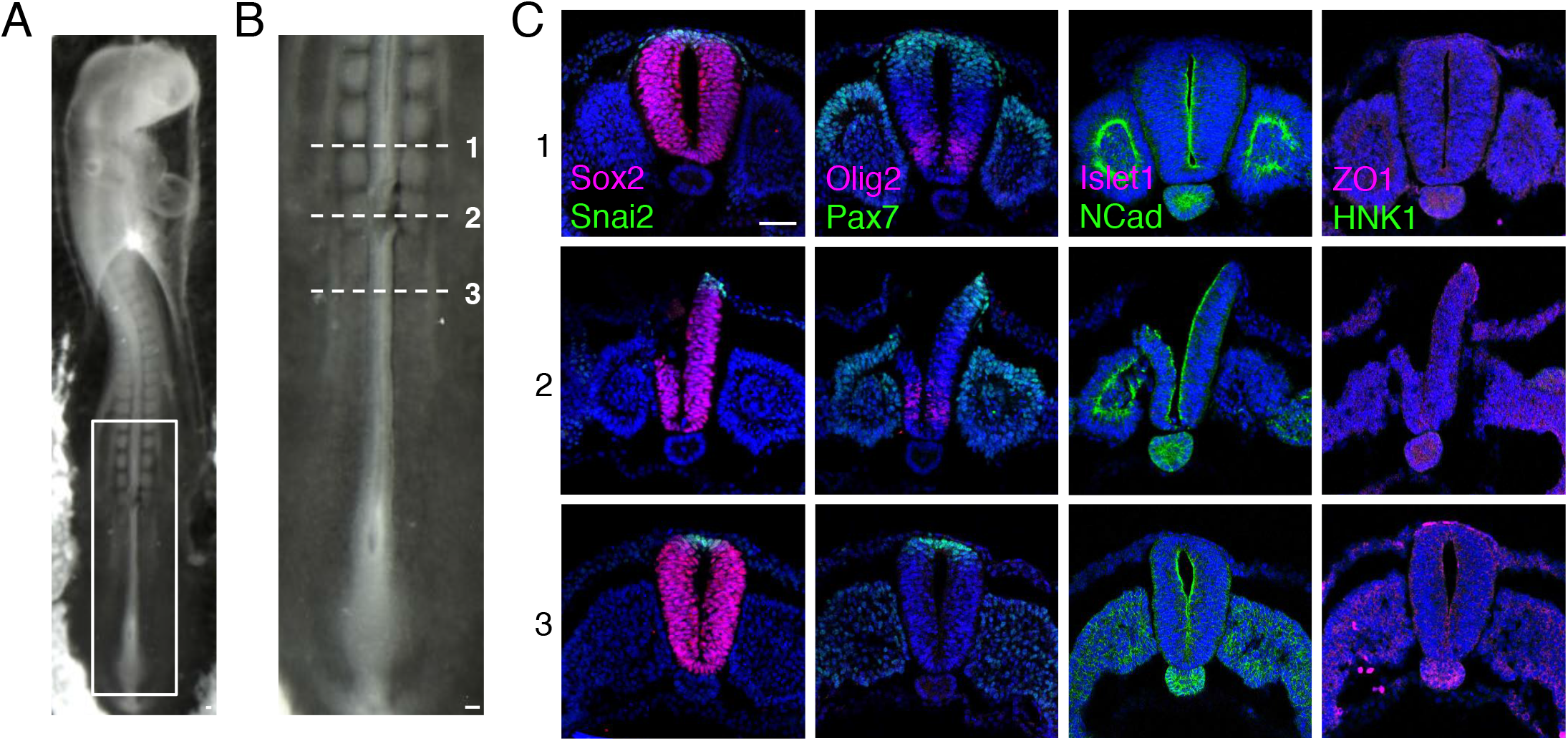
Dorsal region of the chicken neural tube removal prior to grafting. (**A**) E2 (10-15 somite stage, HH 10/11+) chicken embryo following removal of the dorsal part of neural tube. (**B**) High magnification of the newly formed somite region, where the dorsal part of the neural tube was surgically removed prior to grafting. (**C**) Transverse sections of chicken neural tube subjected to IF for SOX2, SNAI2, PAX7, OLIG2, ISLET1, NCAD, ZO1 and HNK1 in the region anterior to the removal site (1), the removal region (2), and the region caudal to the site (3). The neural tube region removed is only composed of neural crest and neural progenitors expressing respectively SNAI2 and PAX7 and is similar to D6 human spinal cord neural rosettes. At least three chicken embryos and three sections were analysed for each marker combination. Scale bar = 50μm.

One possibility is that a community effect (Gurdon 1988) operates within grafted rosettes, which overcomes differentiation signals provided by host tissues. We therefore analysed separately human cells which were directly incorporated into the chick neuroepithelium, either in small groups or as single cells to determine if these differentiated more rapidly (Fig. 4B’). However, no differences were found between contiguous rosettes and incorporated human cell groups (Figs. 4D-G), and in single cells a slight tendency for reduced progenitor marker expression and increased incidence of P27 was not statistically significant.

Moreover, no ISLET-1 positive cells were found in single cells. This reinforces the conclusion that the pace of human neural differentiation in these rosettes is directed by cell intrinsic factors.

Analysis of neural crest differentiation in grafted rosettes after 2 days further confirmed the asynchrony between graft and host cell differentiation progression. SNAI2-expressing neural crest cells where still found within grafted rosettes, while delamination of host neural crest was now complete, as indicated by absence of Snail2 positive cells in the chicken dorsal neural tube (Fig. 5A). Moreover, expression of migratory neural crest markers HNK1 and TFAP2α in the rosette periphery further aligned human rosette differentiation in vivo with that observed in vitro (Figs. 5B, C and S2). As single cells, integrated human cells were more likely to express markers of migrating neural crest cells HNK1 (Fig. 5B) and TFAP2α (Fig. 5C) exhibiting neural crest differentiation progression similar to that found in *in vitro* conditions (Fig. S2).

**Figure 5.**
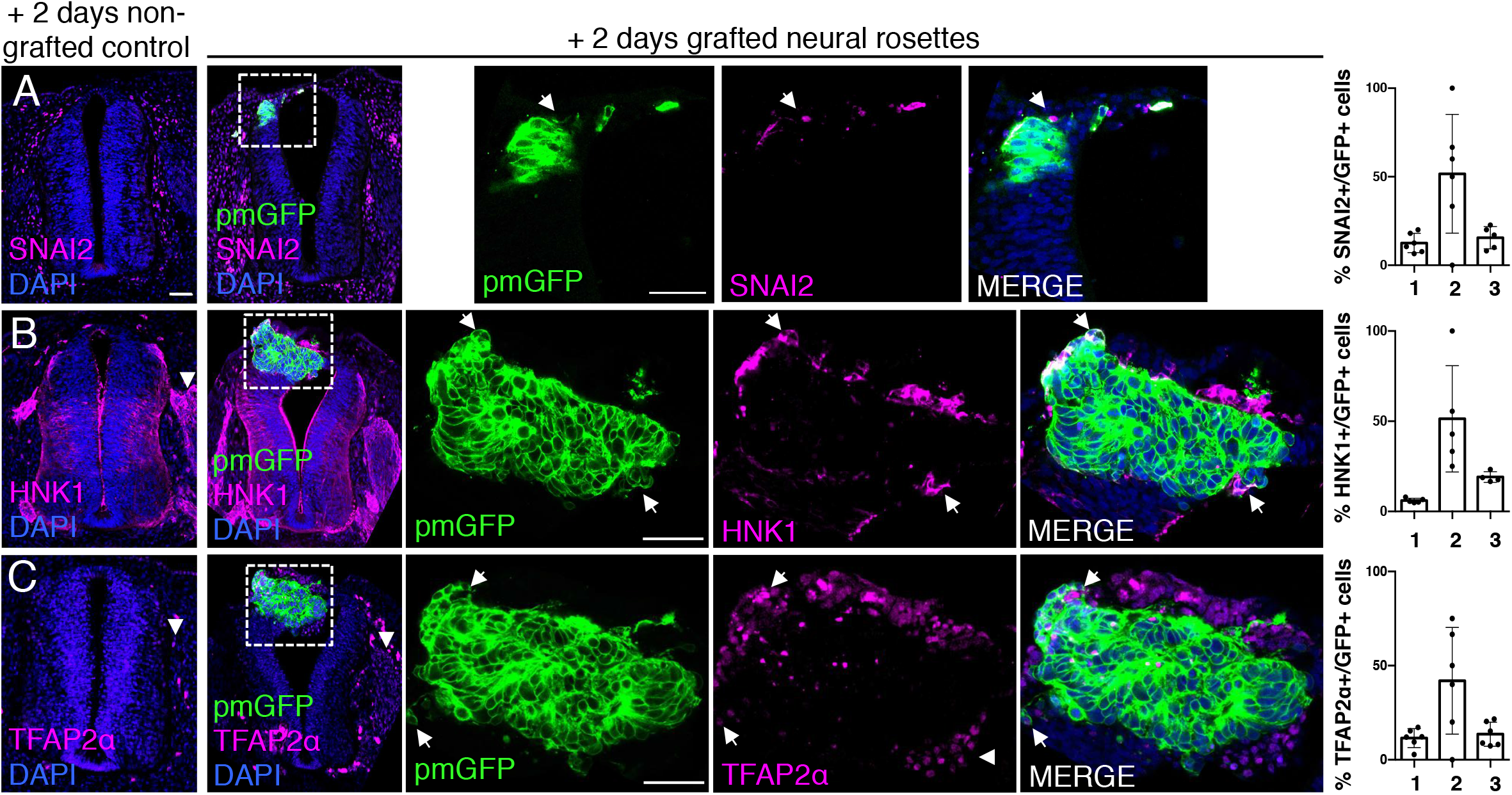
Homo-chronically transplanted hiPSC-derived neural rosettes exhibit cell autonomous neural crest differentiation in the chicken embryonic environment. (**A-C**) IF to detect key neural crest marker proteins in control (non-grafted) and grafted embryos after 2 days incubation (**A**) SNAIL2, (**B**) HNK1, (**C**) TFAP2α (arrowheads indicate examples of candidate protein expressing cells in the chicken host tissue; arrows indicate examples of candidate protein and GFP co-expressing cells, note most cells at graft periphery, dashed white boxes indicate area of magnification in next adjacent image). Three transverse sections from each of at least three grafted chicken embryos were analyzed for each marker and the proportions of SNAIL2/GFP +ve, HNK1/GFP +ve and TFAP2α /GFP +ve cells in cell configurations 1, 2 or 3 were quantified, each dot represents an histological section analyzed, note the small number of isolated cells (group 2) underlies lack of significant difference between groups (see Materials and Methods), data analyzed with Mann-Whitney U test. Errors bars are ± SD. Scale bar = 50μm.

**Figure S4.**
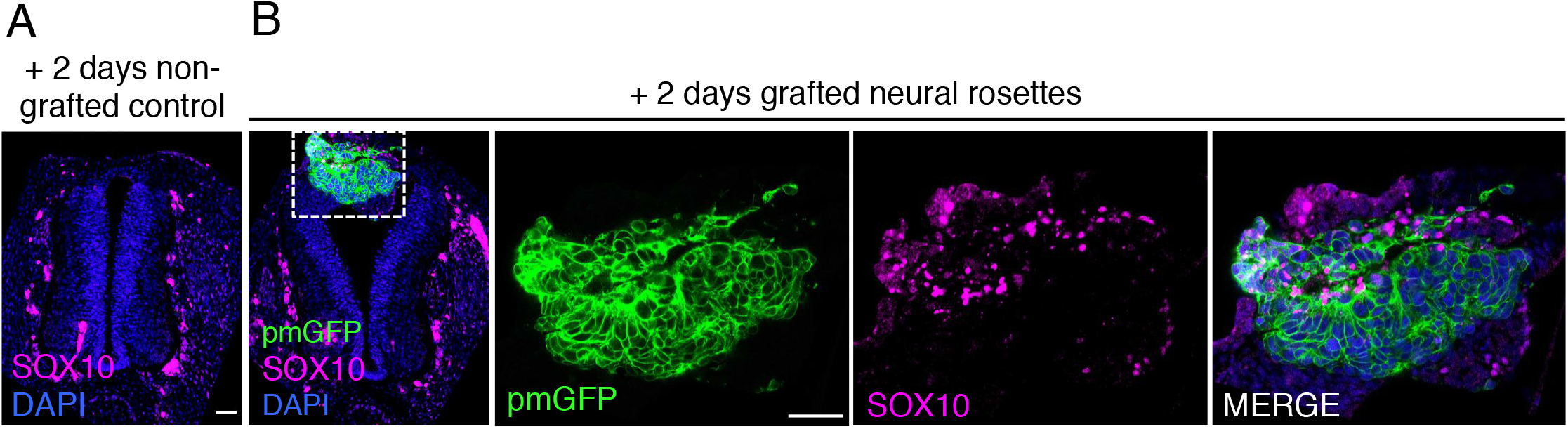
Transplanted human dorsal spinal cord rosettes express SOX10, a specific marker of differentiated neural crest 2 days after grafting. IF to detect expression of the neural crest marker **(A)** SOX10 in control sham operated, and **(B)** grafted chicken embryo following 2 days incubation (dashed white box indicates area of magnification in next adjacent image). One chicken embryo and three sections were analysed. Scale bar = 50μm.

Intriguingly, despite this apparent lack of influence of the chick host on grafted human cells, we found some evidence that human cells altered chick neural crest differentiation. At this stage, chicken neural crest cells have normally delaminated and migrated to the dorsal root ganglia (DRGs) (Kalcheim 1999). However, close to integrated human cells chicken neural crest expressed early migration phase markers HNK1 (Fig. 5B), TFAP2α (Fig. 5C) and Sox10 (Fig. S4) suggesting that the presence of human cells delays progression of the host neural crest differentiation program.

Together these findings indicate that after two days, differentiation in grafted human spinal cord rosettes progresses at the same pace as in vitro despite exposure to the more rapidly differentiating environment of the chicken host embryo.

### Longer term analysis reveals impaired differentiation in grafted human neural rosettes

To address whether the differentiation pace and trajectory in grafted human rosettes continues in synchrony with that in vitro, analysis was further undertaken five days after grafting. At this time point all human cells were incorporated within the host neuroepithelium, either as cell groups or single cells (Figs. 6A-6B’) and the majority of cells remained dorsal neural progenitors expressing SOX2 and PAX7 (Figs. 6C,D). Importantly, neuronal differentiation now continued (Fig. 6E-G) indicated by the increase of P27 and ISLET1 positive human cells. However, the proportion of cells expressing P27 in transplanted rosettes decreased between day 2 and day 5 and was also significantly reduced compared with their in vitro counterparts, where the proportion of P27 cells increased in this time frame (Fig. 6G). The neural marker ISLET1 was first detected five days after grafting, both in vivo and in vitro (equivalent to D11 rosettes) (Fig. 6G, S2). However, the rate of ISLET1 positive cells was significantly reduced in transplanted rosettes compared with their in vitro counterparts (Fig. 6G). Moreover, the neural progenitor cell population continued to expand in vivo to greater extent than in vitro, as indicated by higher proportion of SOX2 and PAX7 expressing cells in transplanted rosettes (Figs. 6G, S2). Together, these results indicate that, the rate of neurogenesis gradually attenuates in transplanted rosettes while the neural progenitor population expands.

**Figure 6.**
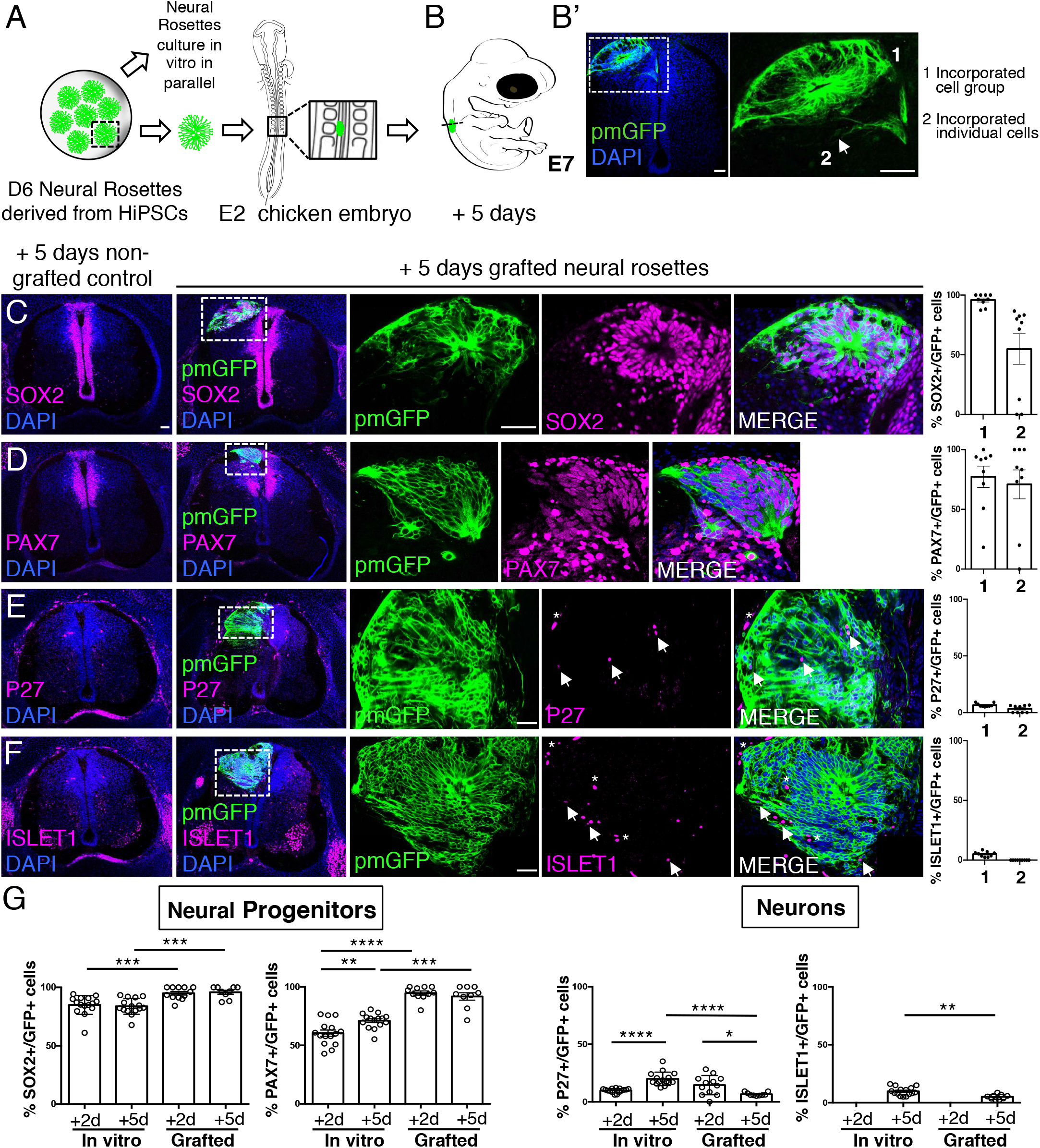
Longer term analysis reveals impaired neuronal differentiation and cell type specification in grafted human neural rosettes. (**A**) Schematic of human dorsal spinal cord neural rosette transplantation protocol and (**B-B’**) transplanted cell configurations 5 days after transplantation; (**C-F**) IF to detect key marker proteins in control (non-grafted) and grafted chicken embryos after 5 days incubation, neural progenitor markers (**C**) SOX2, and (**D**) PAX7, post-mitotic neuronal (**E**) P27 and dorsal interneuron type 3, dI3s, (**F**) ISLET1 markers (arrows indicate examples of P27 and GFP co-expressing cells, dashed white boxes indicate area of magnification in next adjacent image, asterisks indicate examples of host blood cells). Three transverse sections from each of at least three grafted chicken embryos were analyzed for each marker and the proportions of SOX2/GFP +ve, PAX7/GFP +ve, P27 /GFP +ve and ISLET1/GFP +ve cells in configurations 1 or 2 were quantified, each dot represents an histological section analyzed, no significant differences were found; (**G**) Comparison of proportion of ChiPS4-pmGFP derived neural progenitors and neurons expressing SOX2, PAX7, P27 and ISLET1 in rosettes cultured in vitro in parallel with those transplanted into chicken embryos, assessed after 2 and 5 days, each dot represents a single rosette (5 rosettes sampled from each to 3 independent differentiations), see Fig S2 and proportions of these same marker proteins in transplanted rosettes after 2 and 5 days in chicken embryonic environment, each dot represents data from one section, with 3 sections from each of 3 grafted chick embryos, analysis Mann-Whitney U test. Errors bars are ± SD. p values (see methods). Scale bar = 50μm.

Furthermore, day 5 transplanted rosettes lacked the neural crest markers assessed (SNAI2, HNK1 and TFAP2α), but these were detected in rosettes cultured in parallel in vitro (Figs. 7A, B, C and S2). This indicates that this later chicken embryo environment is no longer conducive to human neural crest differentiation. However, we note that human neural crest cells were still generated in one chimera excluded due to open neural tube defects (Fig. S5), suggesting that human neural rosettes retain the potential for such differentiation in an appropriate environment.

**Figure 7.**
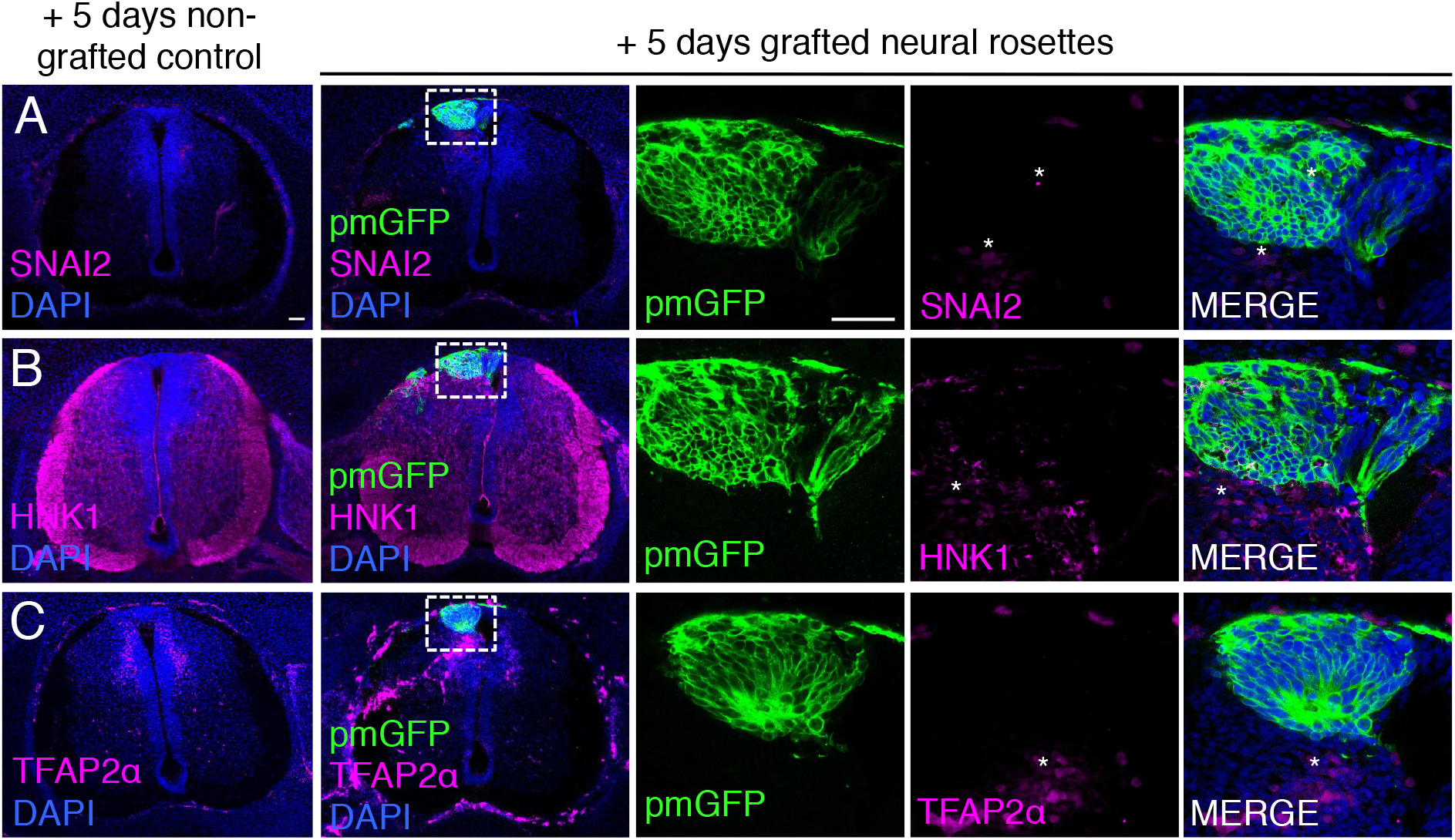
Longer term analysis reveals no neural crest differentiation in grafted human neural rosettes. (**A-C**) IF to detect key neural crest marker proteins in control (non-grafted) and grafted chicken embryos after 5 days incubation reveals no expression of neural crest progenitor markers in any cell configurations (**A**) SNAI2 (1: 0/860 GFP+ cells and 2: 0/70 GFP+ cells), (**B**) differentiated neural crest cells HNK1 (1: 0/956 GFP+ cells and 2: 0/57 GFP+ cells) and (**C**) TFAP2α (1: 0/1175 GFP+ cells and 2: 0/66 GFP+ cells) (asterisks indicate host blood cells). At least three chicken embryos and three cross-sections were analyzed for each marker. Scale bar = 50μm.

**Figure S5.**
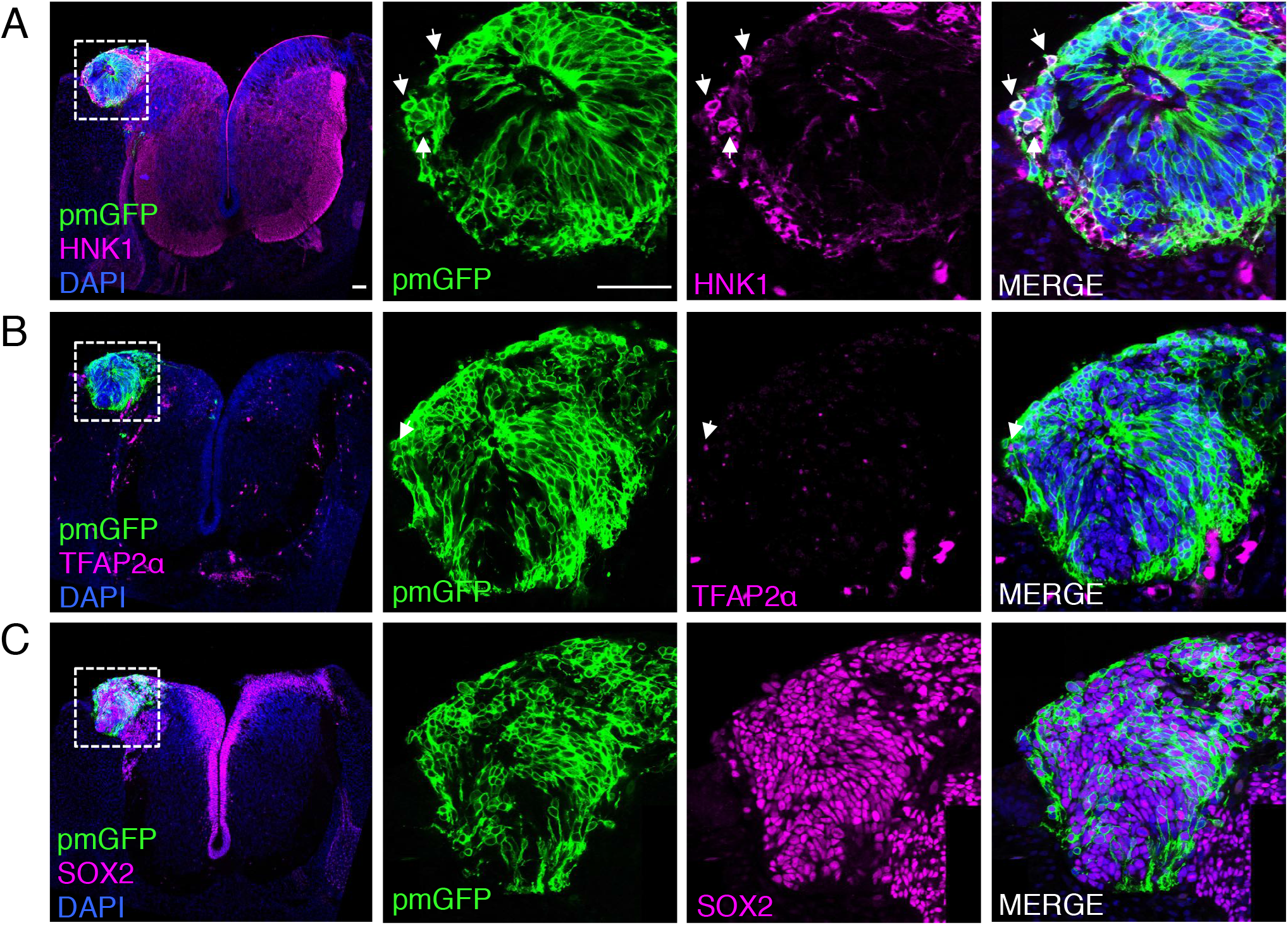
Ectopically positioned human dorsal spinal cord rosette reveals retention of ability to differentiate into neural crest. Chicken open neural tube with a human dorsal spinal cord rosette following transplantation 5 days previously, analysed with IF to detect expression of differentiated neural crest cell markers **(A)** HNK1, **(B)** TFAP2α and (**C)** pan-neural marker SOX2 (arrows indicate examples of candidate proteins and GFP co-expressing cells, dashed white boxes indicate area of magnification in next adjacent image). One chicken embryo and three sections were analysed. Scale bar = 50μm.

Overall, longer term analysis of differentiation in transplanted human rosettes revealed that rather than proceeding more rapidly in the chicken embryo, even the intrinsic differentiation pace and trajectories of human dorsal neural progenitors and neural crest are ultimately not sustained and cells in the human rosettes stall in the expansive neural progenitor phase.

## Discussion

This study presents a novel in vitro human neural rosette assay that recapitulates the temporal sequence of dorsal spinal cord differentiation observed in vertebrate embryos. Importantly, this differentiation programme progresses more rapidly in vitro than in the human embryonic spinal cord. It may therefore capture the cell intrinsic differentiation pace of this cell population, while lacking signals present in the embryo that influence the normal duration of steps in spinal cord differentiation. The initial retention of this in vitro differentiation pace in rosettes challenged by transplantation into the faster differentiating environment of the chicken embryonic spinal cord, further suggests that this reflects a cell intrinsic species-specific differentiation pace. However, analysis of transplanted rosettes after a longer period revealed that not only do these cells fail to respond to differentiation cues in the chick embryonic environment, but also no longer follow their intrinsic differentiation programme, and persist instead in the expansive neural progenitor cell state.

In vitro differentiation of human pluripotent cell derived neuromesodermal progenitors in minimal medium containing low level retinoic acid (Verrier et al. 2018) was sufficient to generate self-organizing dorsal spinal cord rosettes composed of neural crest and neural progenitors. We demonstrated that such structures reproducibly differentiate in a specific temporal sequence which follows that observed in the embryonic dorsal spinal cord (Fig. 3I), with neural crest differentiating first, then dorsal-interneurons (dI1/3/5s) and finally glial cells (Duband, Dady, and Fleury 2015, Jiang et al. 2009, Andrews et al. 2017, Deneen et al. 2006). One peculiarity in this assay, the lack of dI2/4s interneurons, may be explained by a requirement for more nuanced exposure to BMPs (Andrews et al. 2017, Gupta et al. 2018, Duval et al. 2019), which, although endogenously expressed in this assay, are not exogenously provided in this protocol (Verrier et al. 2018).

Two lines of evidence indicate that these human dorsal spinal cord rosettes exhibit a cell intrinsic differentiation pace. First, the timing of neural progenitor differentiation was found to be more rapid in vitro than in the human embryo. This was indicated by the earlier appearance of ISLET1+/HB9-expressing neurons and of NFIA expressing gliogenic progenitors in neural rosettes than in the human embryonic dorsal spinal cord; indeed, the period from migratory neural crest emergence to gliogenesis onset was ∼1.5 weeks faster in vitro than in the human embryo (Fig 3I). This detailed comparison of the timing of the appearance of specific cell types in the human embryo with in vitro data may help to inform modelling approaches which largely rely on morphological changes to align differentiation tempo (Workman et al. 2013). This discrepancy in differentiation timing supports the idea that in vitro conditions represent a differentiation pace that reflects cell intrinsic constraints. Secondly, homo-chronic transplantation of these rosettes into the chicken neural tube, which has a more rapidly progressing differentiation tempo, did not accelerate rosette differentiation (even in individual human cells incorporated within the chicken neuroepithelium), further suggesting that rosette differentiation pace reflects species-specific cell intrinsic constraints. This conclusion is consistent with the distinct differentiation pace manifest in vitro in human and in mouse ventral neural progenitors cultured in similar conditions, which have been shown to reflect species-specific differences in cell cycle length and general protein stability (Rayon et al. 2020).

In this context, it is informative that our experiments demonstrate that the pace of human neural differentiation in vitro is faster than that observed in the human embryo. This likely reflects longer time spent in expansive neural progenitor phases in the embryo, where this leads to generation of a much larger structure than a rosette. This raises the interesting possibility that overall differentiation pace, beyond that governed by cell intrinsic factors, might be influenced by and scale with the size of a self-organising cell group. In vertebrate embryos, however, the timing of specific phases of neural differentiation is known to be orchestrated by signals from adjacent tissues, which drive and coordinate axial differentiation: this includes adjacent somites (Diez del Corral, Breitkreuz, and Storey 2002, Diez del Corral et al. 2003), underlying notochord (Placzek et al. 1990) and overlying ectoderm/roof plate (Liem, Tremml, and Jessell 1997, Lee, Dietrich, and Jessell 2000). Clearly, the localised and temporally controlled characteristics of such regulatory influences are lacking this in vitro assay. The importance of timely exposure to extrinsic cues is further underscored by the cell intrinsic behaviour of transplanted human rosette cells, despite exposure to signals from an appropriate extra-cellular environment. One possibility is that the slower human cell cycle (Rayon et al. 2020) contributes to this lack of response to endogenous chick differentiation cues. Indeed, transcriptional onset of differentiation genes may require exposure to signals in a specific cell cycle phase (Pauklin and Vallier 2013, Pauklin et al. 2016) and involve defined changes in replication timing at the early G1 timing decision point (Dimitrova and Gilbert 1999, Hiratani et al. 2008). Perhaps related to this is the finding that while human and mouse neural progenitor cells similarly transduce exogenously provided signalling molecule sonic hedgehog, human cells are slower to transcribe *OLIG2* and *NKX6*.*1*, key downstream ventral patterning genes (Rayon et al. 2020).

Previous studies have used xenografting assays to evaluate the behaviour of human progenitors generated *in vitro* (Valensi-Kurtz et al. 2010, Frith et al. 2018) or for potential long-term regenerative therapy purposes using either neural progenitor cells or differentiated neurons, e.g. (Kumamaru et al. 2018, Kumamaru et al. 2019, Linaro et al. 2019). Here we have focused on the differentiation potential of human neural progenitors in an ectopic cellular environment. We have found that these cells only initially follow the intrinsic differentiation programme that they exhibit in vitro. Five days following rosette transplantation neuronal differentiation declines compared to in vitro conditions, neural crest differentiation ceases and cells retain dorsal neural progenitor gene expression. This loss of differentiation progression reveals a limitation of the cell intrinsic neural differentiation programme and underscores the requirement for appropriate extrinsic inputs to maintain differentiation trajectory. Indeed, this suggests that differentiation signals are soon lacking in the chick embryonic environment but are provisioned sufficiently in minimal culture conditions: this might include exogenously provided retinoic acid. In addition, we cannot rule out inhibitory signals acting on grafted human neural cells. Most striking was the continued expansion of these no longer differentiating transplanted neuroepithelial cells; a behaviour reminiscent of neural cancers such as Glioblastoma that share a similar neural progenitor gene expression profile (Agnihotri et al. 2013, Hassn Mesrati et al. 2020). In addition to providing novel insight into the regulation of human neural differentiation, this assay may therefore also offer an opportunity to study cancer origins as well as inform cell-based therapeutic approaches to neural injury and disease.

## Materials and Methods

### hESC and hiPSCs culture and differentiation

hESC (H9 or AI03e-DCX-YFP H1 line) and hiPSCs (ChIPS4, Cellartis AB) were maintained as feeder-free cultures in DEF-based medium supplemented with bFGF (30ng/mL, Peprotech, Cat. No. 100-18B) and Noggin (10ng/ml, Peprotech, Cat. No. 120-10C) on Geltrex matrix coated plates (10μg/cm^2^, Life Technologies, Cat. No. A1413302), and enzymatically passaged using TryPLselect (Thermofisher, Cat. No. 12563011). For PSC passaging, the medium was complemented by addition of the Rho kinase inhibitor Y-27632 (10mM, Tocris, Cat. No. 1254). HPSC quality control and passage number information are provided as metadata.

For differentiation assays, PSC were plated on Geltrex matrix at 40000cells/cm^2^ and shifted to N2B27 medium after 24h. Cells were then differentiated for 3 days (D) in N2B27 supplemented with 3 µM Chiron99021 (Tocris, Cat. No. 4953) and 20ng/ml bFGF (PeproTech, Cat. No. 100-18B), in the presence of 50ng/mL Noggin (Peprotech, Cat. No. 120-10C) and 10ng/mL SB431542 (Tocris, Cat. No. 1614) from D2 to D4 to generate NMPs, see (Verrier et al. 2018). For further differentiation, NMPs were passaged using PBS-EDTA 0.5mM and seeded back at controlled density (10^5^ cells/cm^2^) on Geltrex (20μg/cm^2^, Life Technologies, Cat. No. A1413302) in the presence of Y-27632 (10mM, Tocris, Cat. No. 1254). Cells were then cultured in N2B27 containing 100nM all-trans-Retinoic Acid (SIGMA, Cat. No. R2625) for the indicated time to obtain and differentiate neural progenitors. First human neural rosettes were observed from D06 with this regime. All work with hESCs was undertaken in accordance with the UK Stem Cell Bank steering committee (license numbers SCSC14-28 and SCSC14-29).

### Immunofluorescence

Primary antibodies for immunofluorescence were obtained commercially for SOX2 (1/200; Millipore), ZO1 (1/500; Invitrogen), NCAD (1/500; SIGMA), TUJ1 (1/1000; SIGMA (Rabbit) or BioLegend (Mouse), OLIG2 (1/200; Millipore), LHX2 (1/200, SIGMA), TLX3 (1/200, Abcam), ISLET-1 (1/200; Abcam), HB9 (1/10; DHSB), PAX7 (1/10; DHSB), SNAI2 (1/200; BD Biosciences), SOX10 (1/200; BD Biosciences), TFAPα (1/10, Santa Cruz), HNK1 (1/10; DHSB), P27 (1/100, BD Biosciences). Human embryonic tissue fixed in 4% PFA for 2h at 4°C was provided by HDBR tissue bank (project no 200407) and processed for cryo-sectioning using standard procedures. hESCs or iPSCs were fixed in 1% Paraformaldehyde solution overnight at 4°C, permeabilized with 0.1% Trixton X-100 in PBS 1X, subjected to immunofluorescence labeling using appropriate secondary antibodies conjugated to Alexa-fluor 488, 568 or 594 (Life Technologies), and processed for DAPI staining to visualize cell nuclei. Fluorescent images were captured using a Leica SP8 confocal using a 20X air or 63X NA1.2 APO water immersion objectives.

### Pm-e GFP Plasmid construction for engineering h IPS cells

A plasmid was constructed for engineering hIPSCs with the piggybac transposase system using the aPX1 transposon plasmid (kind gift from Timothy Sanders, University of Chicago), containing the cytomegalovirus (CMV) enhancer, chicken b-actin (CAG) promoter expression and including the Invitrogen Gateway RfA cassette, allowing for phiC31-mediated recombination from Gateway Entry vectors. Finally, the CAG promoter and Gateway cassette (cargo) are flanked by two Inverted Terminal Repeat sequences (ITR, TTAA) allowing the piggyback transposon action. To label the cell membrane, a plasma membrane (pm) associated palmitoylated fluorescent protein was generated by the addition of the 20-amino acid sequence of ratGAP-43 MLCCMRRTKQVEKNDEDQKI to the N-terminus of the monomeric enhanced GFP (eGFP) (K. Svoboda, Addgene plasmid 18696) through sequential PCR amplification to make a pm-eGFP sequence. This pm-eGFP sequence was then inserted into the aPX1 donor vector using the Gateway System^©^ (ThermoFisher).

### HiPSCs transfection

HiPSCs (Cellartis hIPS4 cell line) cells were transfected using the Neon™ transfection System (Life Technologies, Cat. No. MPK10025). Briefly, a mix of aPX1-pm-eGFP construct and piggyback transposase (Yusa et al. 2011), with a final concentration of 1-2 μg/μl, was added to a solution of 10^7^ cells and a single pulse of 1100V with 30 ms duration was delivered via the Neon transfection system^©^. Transfected cells were then plated back on Geltrex coated dishes (10μg/cm2, Life Technologies) and cultured in DEF-based medium supplemented with bFGF (30ng/mL, Peprotech) and Noggin (10ng/ml, Peprotech).

### Chick embryo grafting assay

Fertilized chicken eggs (Gallus gallus domesticus) from a commercial source (Winter Farm, Hertfordshire-Royston, UK) were incubated at 38°C to Hamburger and Hamilton (HH) stage 11-12 (13 to 16 somites) (Hamburger 1951). A polyclonal population of transfected hIPS4-pm-eGFP cells were cultured 6 days from NMPs differentiation, newly formed human neural rosettes were then manually micro-dissected using a sharp glass needle (0.290-0.310 mm) (Vitrolife). After resting in CO_2_ incubator at 37°C for 2 hours, cell aggregates (diameter ≈ 50-100 μm) were gently dissociated and implanted into the dorsal region of one side of the neural tube of stage HH 11-12 chicken embryos. Before implantation, the dorsal region on one side of the neural tube was removed at the level of the most recently formed somite (Figure S3). Embryos were then harvested 2 days or 5 days post-transplantation (Stage HH 23-24 and HH 30-31 respectively). Chicken embryos were fixed in 4% PFA overnight at 4°C, rinsed in PBS and left in PBS/30% sucrose solution overnight at 4°C, then embedded and in agar, frozen and cryo-sectioned at 20 μm and subjected to immunofluorescence processing (as above).

### Cell counts and statistical analysis

At D4 of in vitro differentiation from NMP-like cells, defined fields (200 μm^2^) of neural progenitors forming a neural plate-like shape were selected for quantification. At later time points, human neural rosettes were selected for analysis and quantification by their shape (round with a single lumen) and size (neural rosette diameter, <u>D6</u>: 70-100 μm, average: 84 μm ± 2,7 μm; <u>D10</u>: 125-137 μm, average: 129 μm ± 1,5 μm; <u>D14</u>: 155-195 μm, average: 179 μm ± 2,7 μm). All cell counts are an average percentage of cells positive for each protein of interest compared to total (DAPI+ or GFP+) cells counted within a field of neural progenitors (D4) or a rosette and contacting cells at its periphery. Fifteen fields of neural progenitors (D4) or 15 rosettes (D6 and later) were analysed for each marker (n=15, 5 fields of neural progenitors or rosettes from each of 3 independent differentiation experiments). For transplanted human rosettes at least 3 sections from grafts in 3 different embryos were used to determine the proportion of cells expressing a protein of interest. Transplanted human cells were only counted when cells were positive for GFP, immunoreactivity for protein of interest and DAPI. To avoid counting the same cells in consecutive sections, alternate sections were used for each different immunolabelling assay (4 different immunofluorescent assays/transplanted embryos)(metadata Tables S1-7). Statistical analysis was performed using non-parametric Mann-Whitney U test for none normally distributed data using Prism V8. Results are represented as means +/- SEM or SD as indicated in figures and p-values are indicated with * P<0.05, ** P<0.01, *** P<0.001 and **** P<0.0001.

## Acknowledgements

We are grateful to Aida Rodrigo-Albors and to Teresa Rayon and James Briscoe for critical reading of this manuscript. We also thank Dr Boaz Levi (Allen Institute) and WiCell for the hESC H1 line expressing DCX^Cit/Y^. Human embryonic material was provided by the Joint MRC/Wellcome Trust (grant# MR/R006237/1) Human Developmental Biology Resource (www.hdbr.org, project no 200407). This work was supported by a Wellcome Investigator award WT102817AIA and an enhancement award WT102817/Z/13/A to KGS. The confocal microscope used for imaging was purchased with support from Wellcome Trust Multi-User Equipment grant (WT101468).

---

**Table 1.**
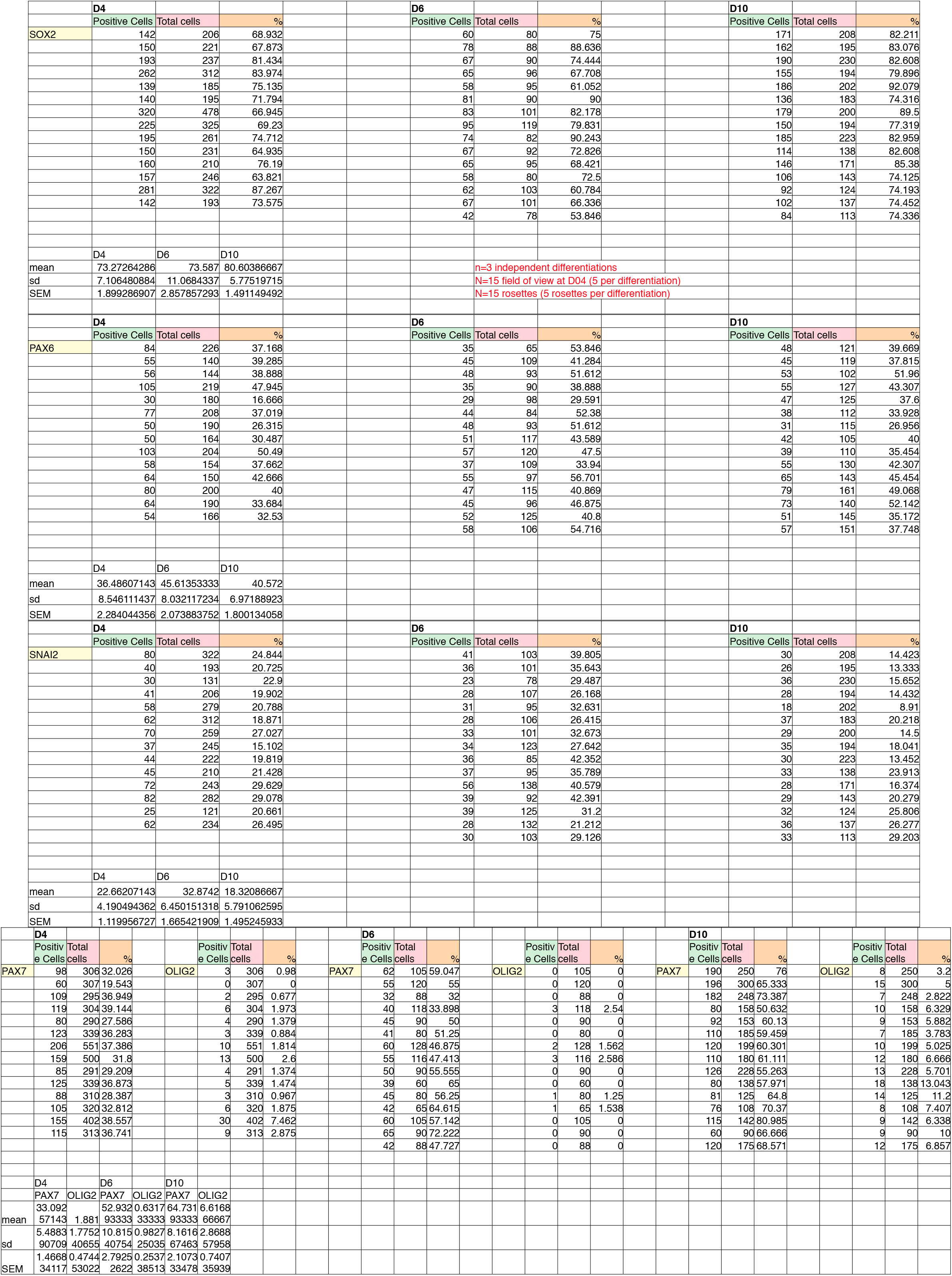
Figure 1 quantification

**Table 2A.**
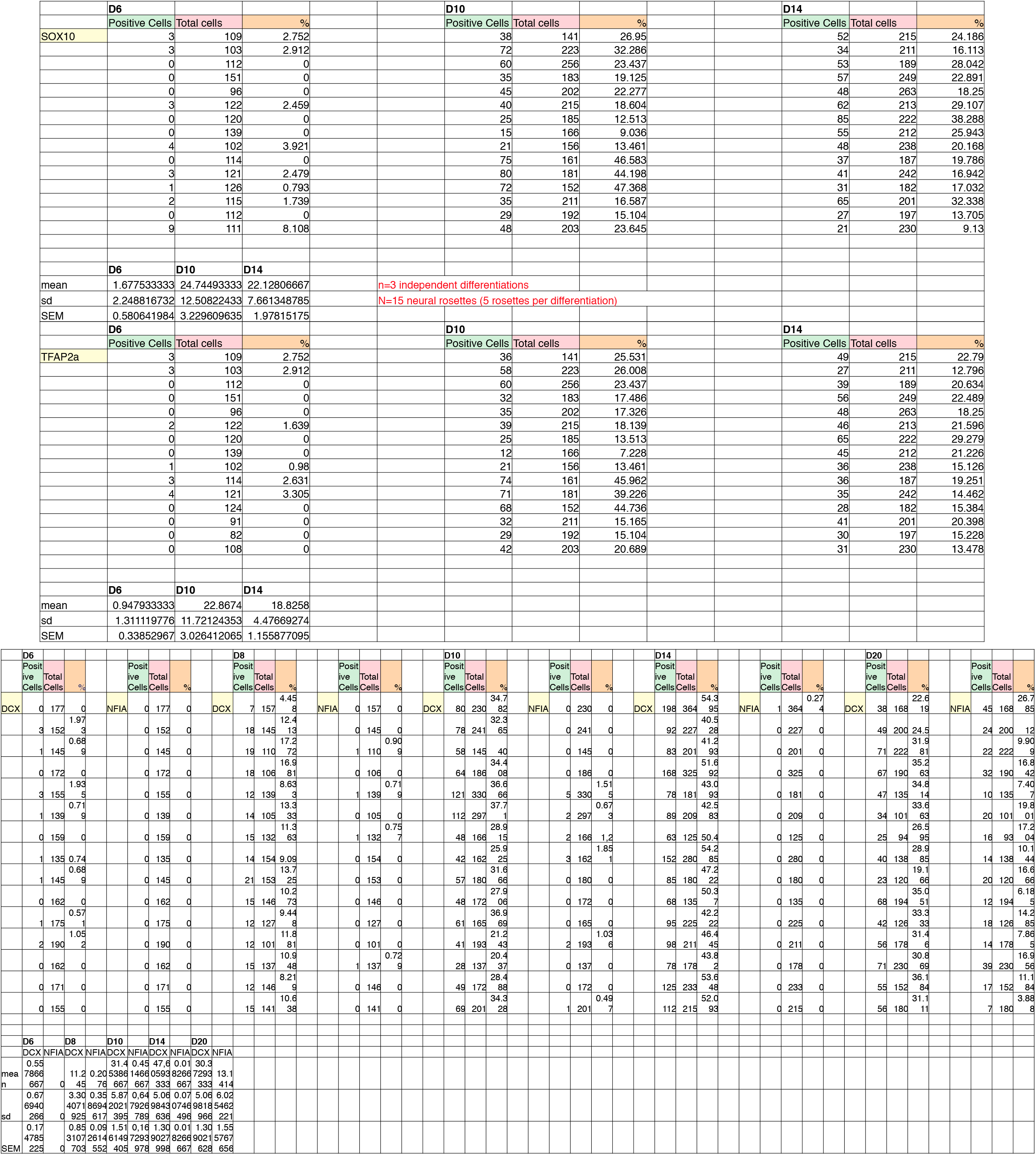
Figure 2 quantification

**Table 2B.**
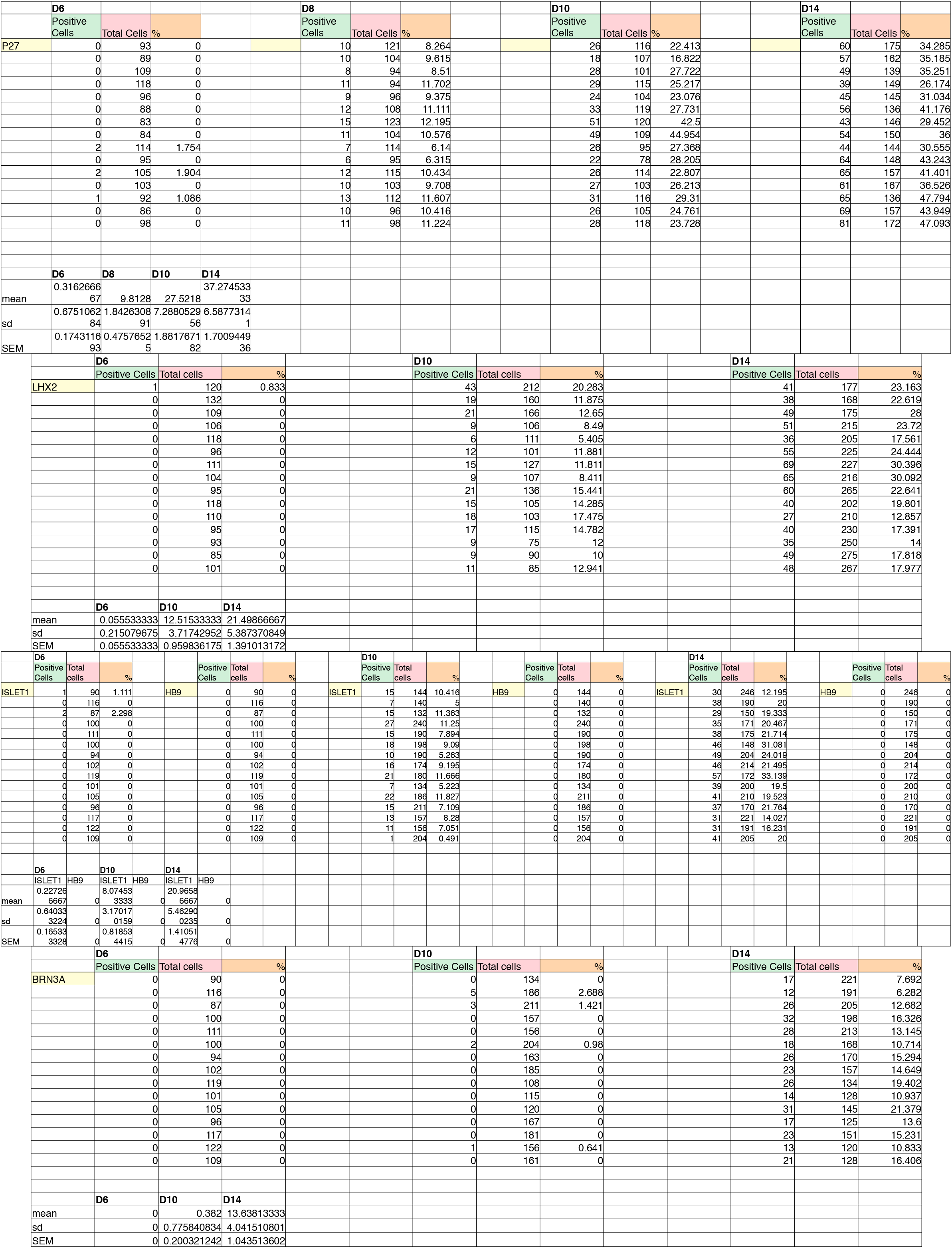
Figure 2 quantification

**Table 3.**
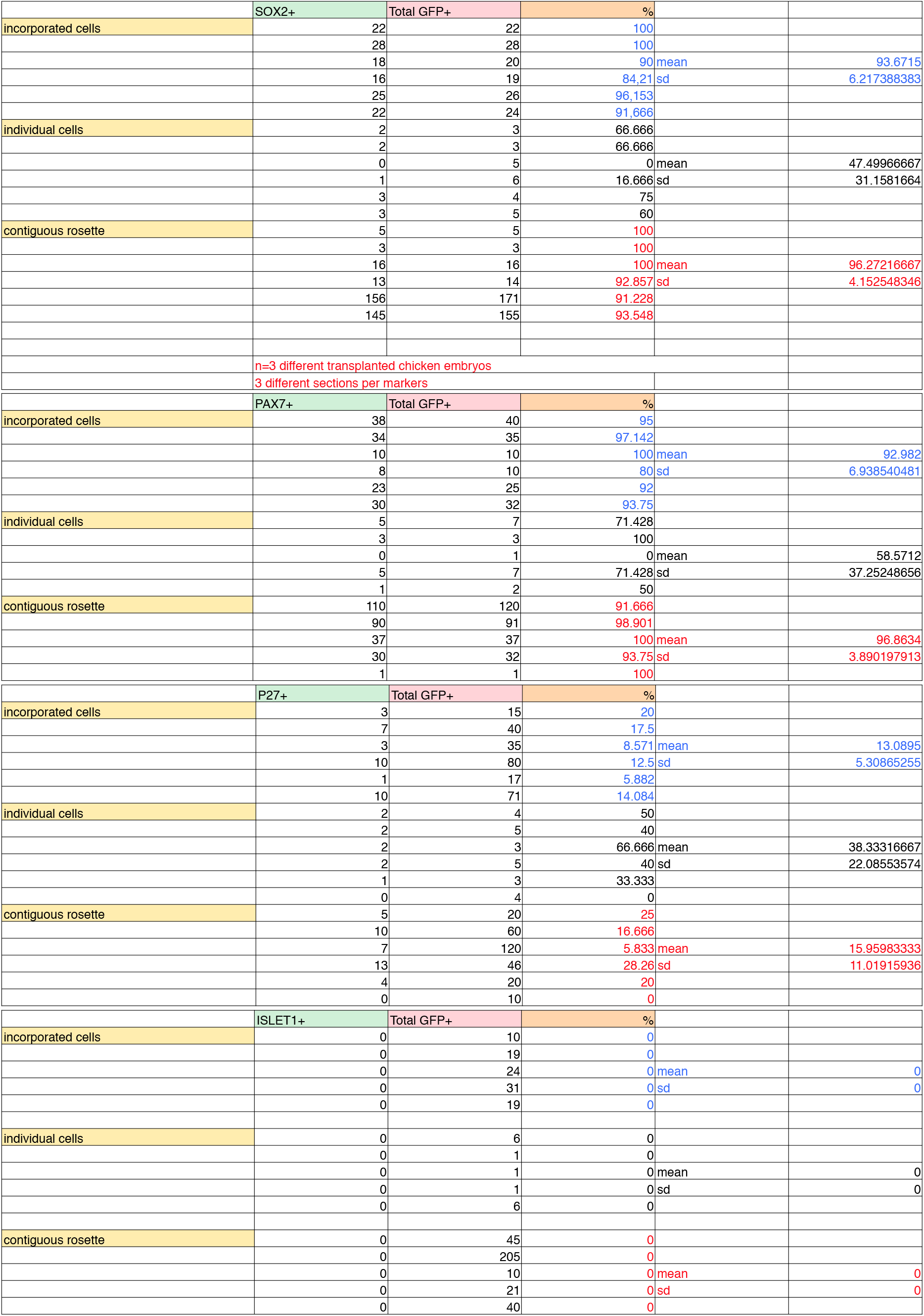
Figure 4 quantification

**Table 4.**
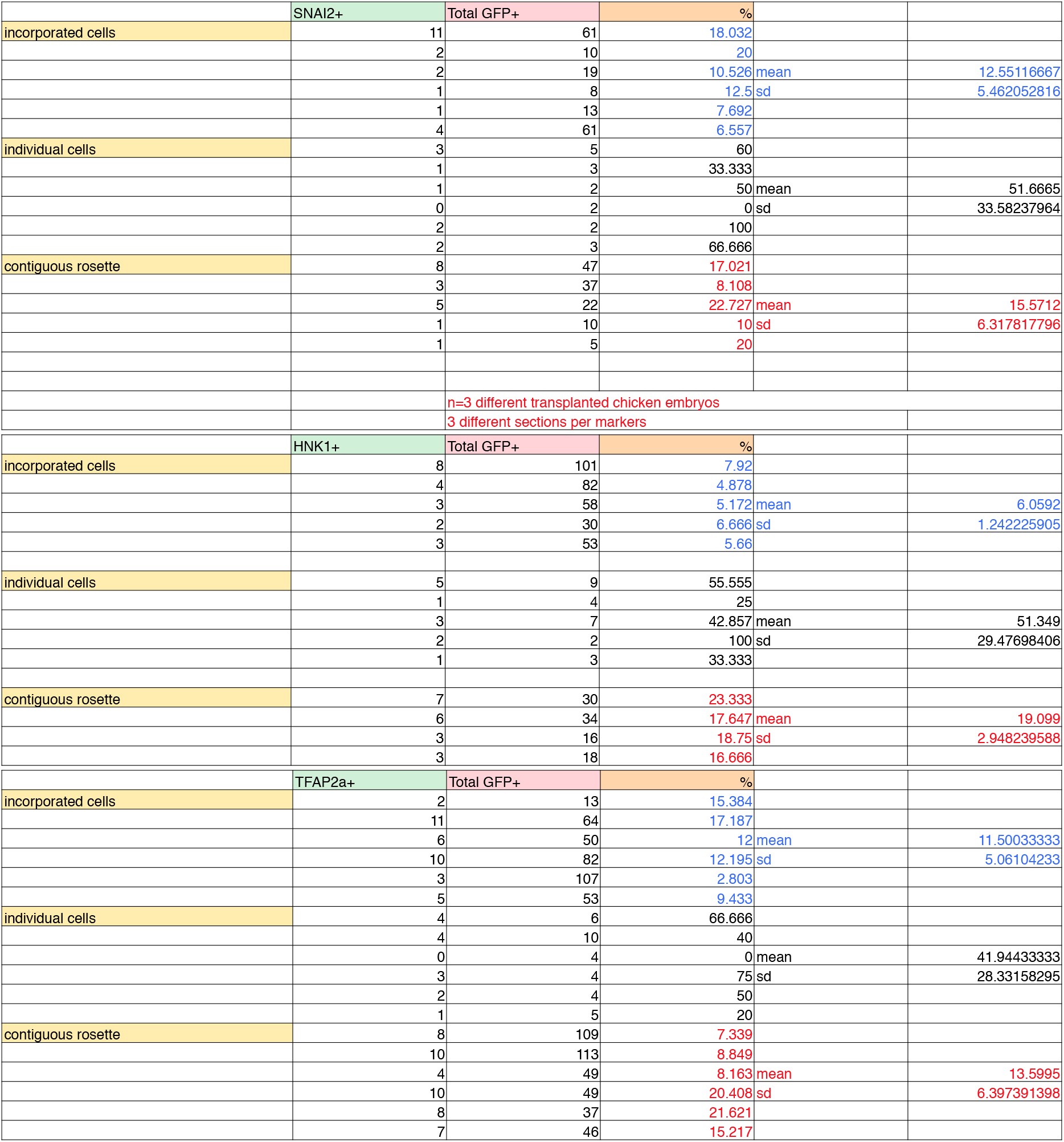
Figure 5 quantification

**Table 5A.**
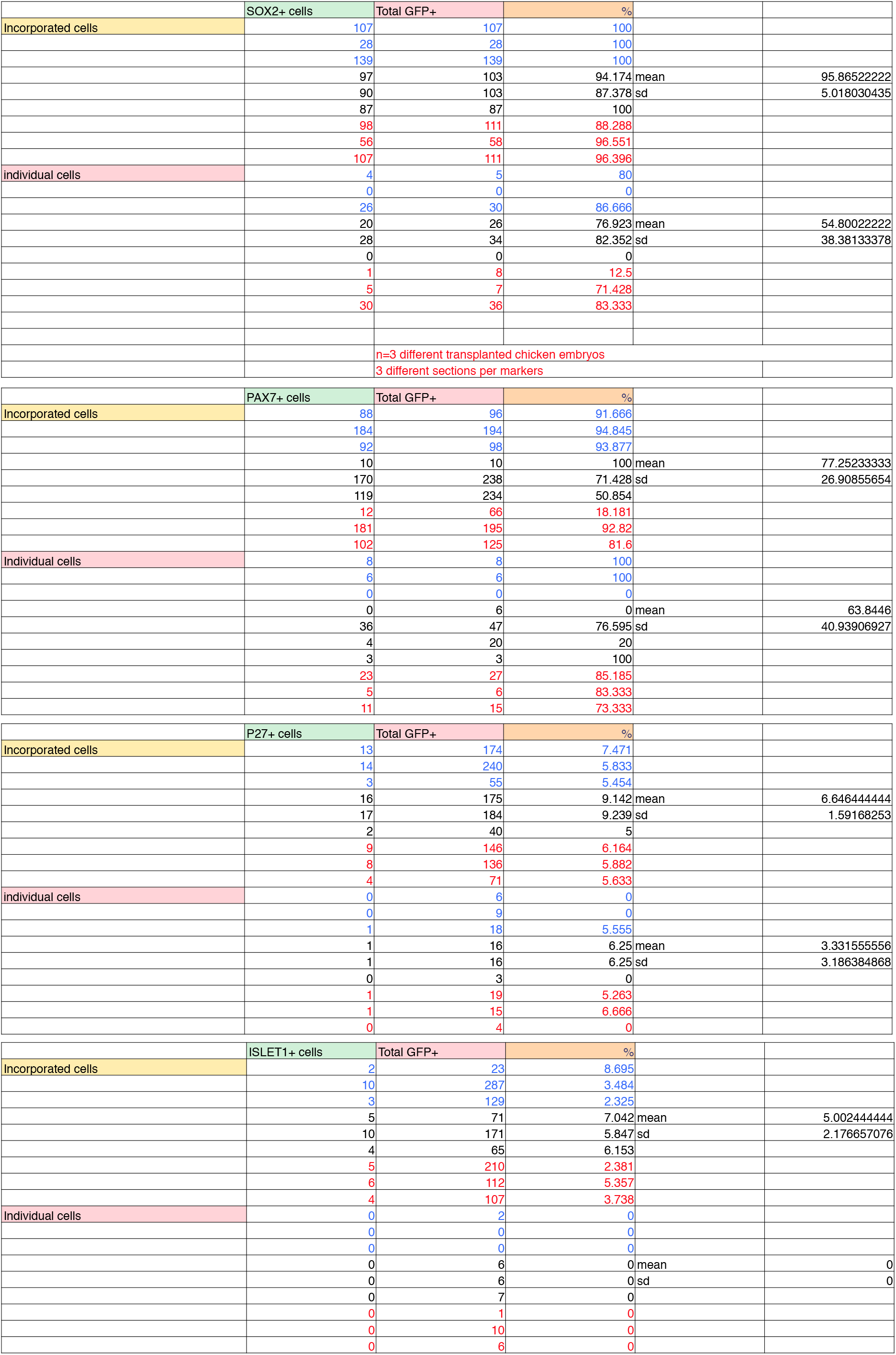
Figure 6 quantification

**Table 5B.**
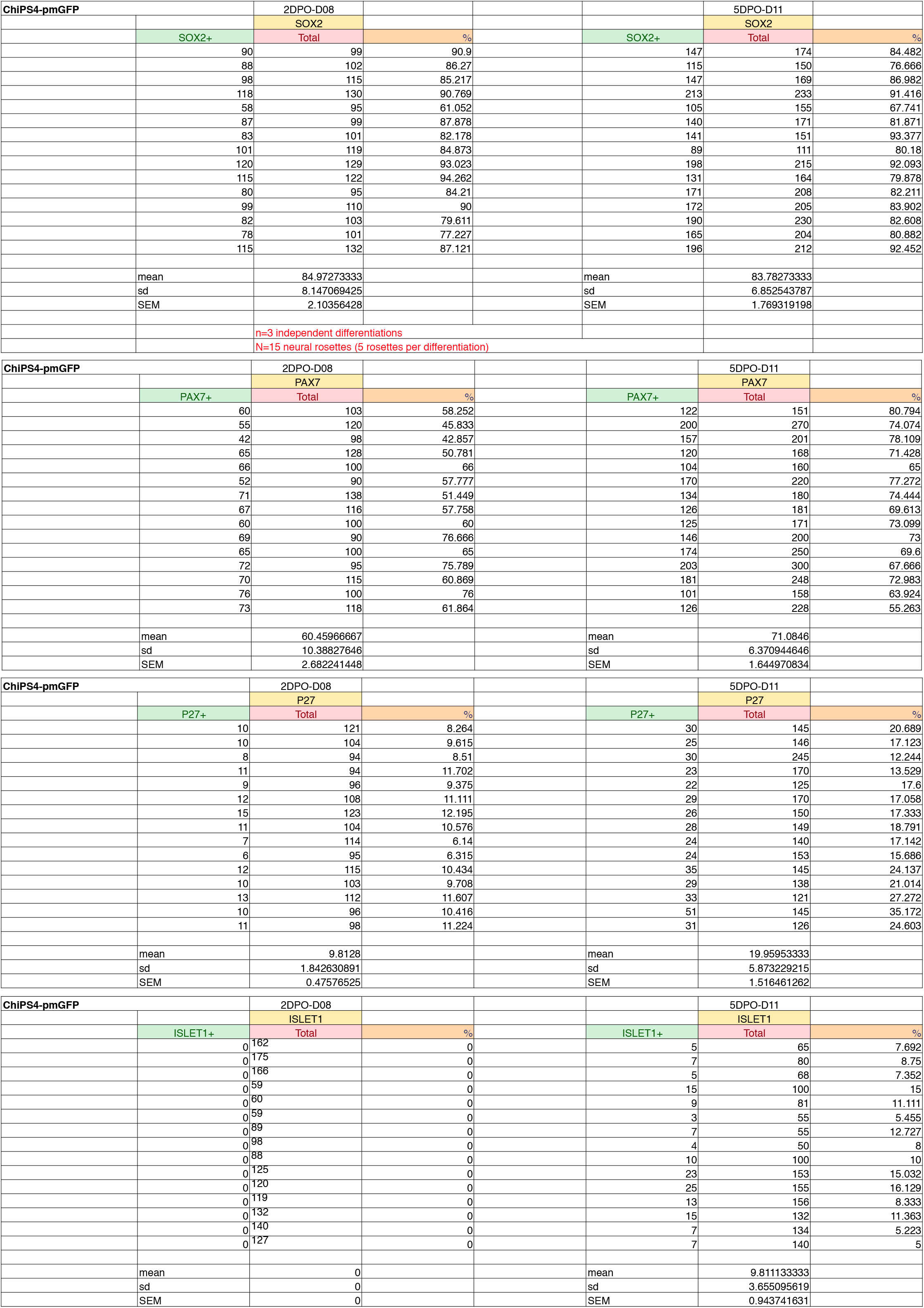
Figure 5 quantification

**Table 6.**
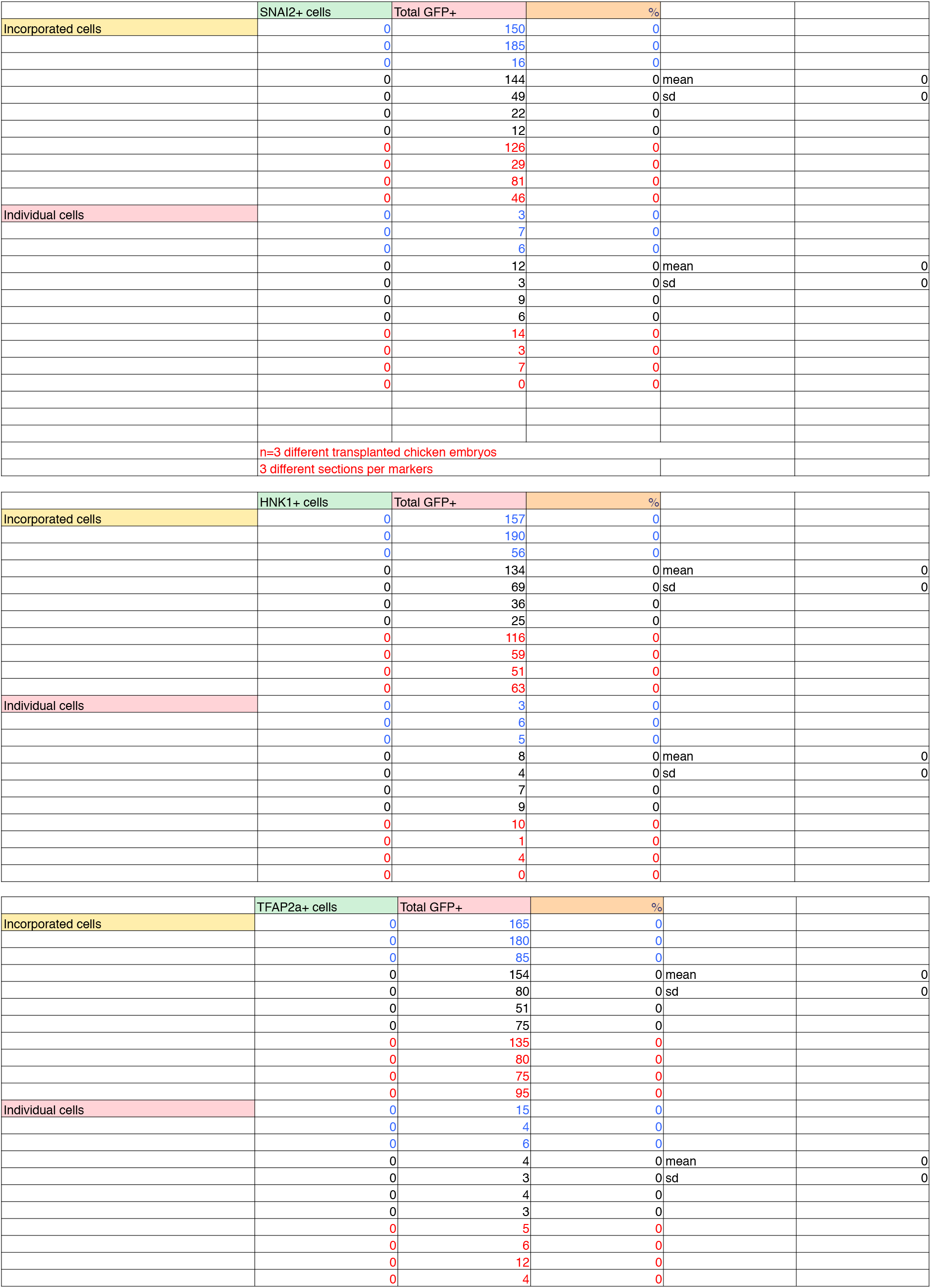
Figure 7 quantification

**Table 7.**
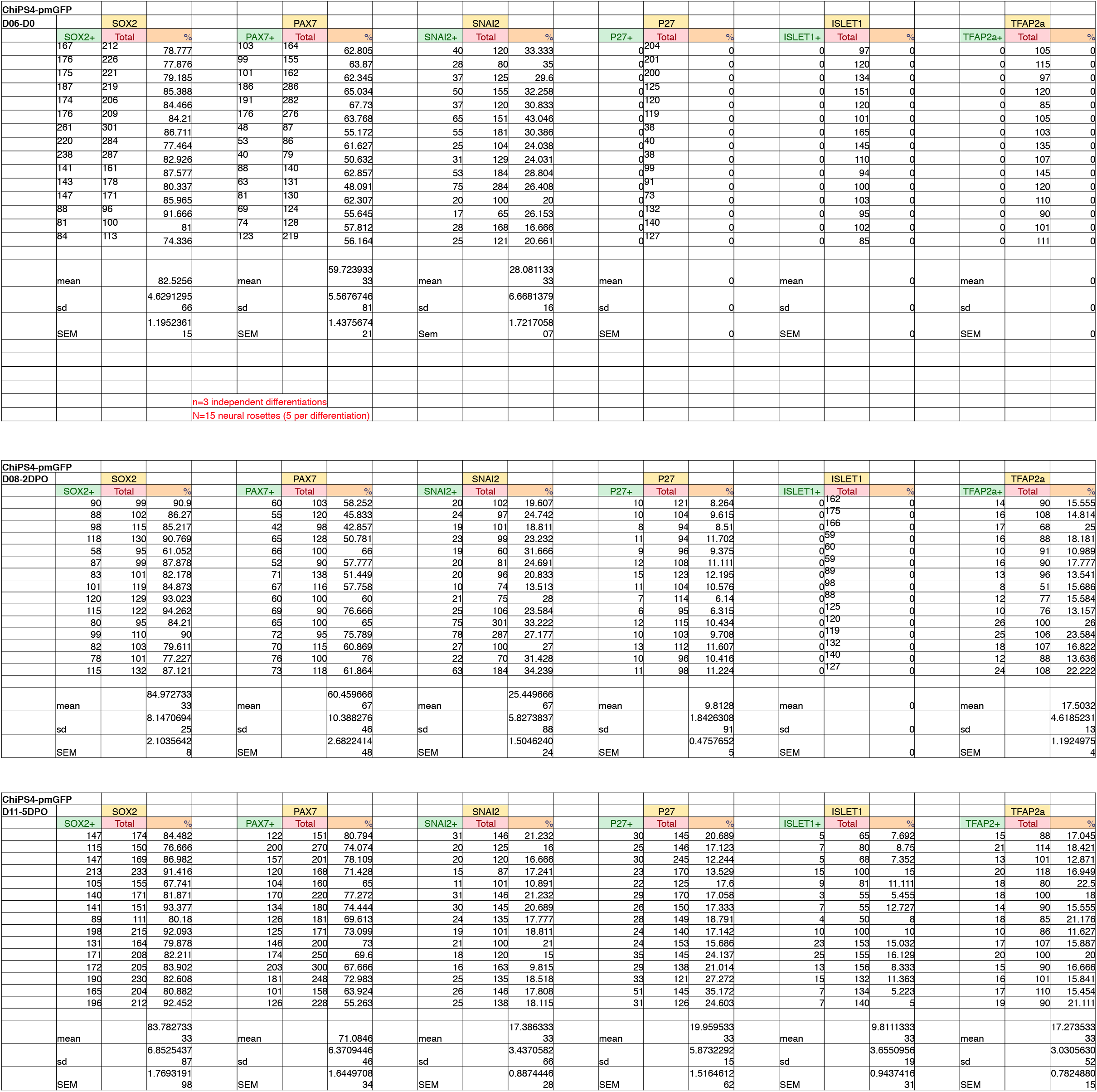
Figure S2 quantification

## Quality control and passage numbers for pluripotent cells used in this study

H9 (WA09) hES cells were purchased from Wicell and were supplied at passage 24. The cells were thawed and cell banks prepared at passage 29. For experiments the cells were used between passage 29 and 39.

H1 (WA01) hES cells were purchased from Wicell and were supplied at passage 30. The cells were thawed and cell banks prepared at passage 34. For experiments the cells were used between passage 34 and 44.

H1-***DOUBLECORTIN*-*CITRINE*** cells were obtained from the Allen Institute and were supplied at passage 63. The cells were thawed and cell banks prepared at passage 66. For experiments the cells were used between passage 66 and 76.

ChiPS4 hiPS cells were purchased from Cellartis AB and were supplied at passage 9. The cells were thawed and cell banks prepared at passage 13. For experiments the cells used between passage 13 and 23.

To make ChiPS4-pmGFP, the cells were transfected with the PiggyBac and transposase constructs at passage 17 and GFP +ve cells selected by fluorescence activated cell sorting at passage 19. The polyclonal cell line was banked at passage 23. For experiments the cells were used between passage 23 and 33.

For quality control purposes, representative lots of each cell bank were thawed and tested for post-thaw viability, and to ensure sterility and absence of mycoplasma contamination. After 2 passages the cell lines were tested to check for the uniform expression of pluripotency markers (Oct4, Sox2, Nanog, SSEA-3, SSEA-4, TRA-1-60 and TRA-1-81) and absence of differentiation markers (SSEA-1, HNF-3 beta, beta-III-tubulin and smooth muscle alpha-actinin) by immunofluorescence.

